# Persistent cell migration emerges from a coupling between protrusion dynamics and polarized trafficking

**DOI:** 10.1101/2021.03.20.436273

**Authors:** Kotryna Vaidžiulytė, Anne-Sophie Macé, Aude Battistella, William Beng, Kristine Schauer, Mathieu Coppey

**Affiliations:** Laboratoire Physico Chimie Curie, Institut Curie, PSL Research University, Sorbonne Université, CNRS UMR168, 75005 Paris, France; Institut Curie, PSL Research University, CNRS UMR144, Cell and Tissue Imaging Facility (PICT-IBiSA), 75005 Paris, France; Cell Biology and Cancer Unit, Institut Curie, PSL Research University, Sorbonne Université, CNRS UMR144, 75005 Paris, France; Faculty of Science and Engineering, Sorbonne Université, 75005 Paris, France; Tumor cell dynamics Unit, Inserm U1279, Gustave Roussy Institute, Université Paris-Saclay, 94800 Villejuif, France

**Keywords:** persistent migration, Golgi, RhoGTPase, optogenetics, polarized trafficking, model, intracellular organization, internal polarity, Cdc42, Rab6

## Abstract

Migrating cells present a variety of paths, from random to highly directional ones. While random movement can be explained by basal intrinsic activity, persistent movement requires stable polarization. Here, we quantitatively address emergence of persistent migration in RPE1 cells over long timescales. By live-cell imaging and dynamic micropatterning, we demonstrate that the Nucleus-Golgi axis aligns with direction of migration leading to efficient cell movement. We show that polarized trafficking is directed towards protrusions with a 20 min delay, and that migration becomes random after disrupting internal cell organization. Eventually, we prove that localized optogenetic Cdc42 activation orients the Nucleus-Golgi axis. Our work suggests that polarized trafficking stabilizes the protrusive activity of the cell, while protrusive activity orients this polarity axis, leading to persistent cell migration. Using a minimal physical model, we show that this feedback is sufficient to recapitulate the quantitative properties of cell migration in the timescale of hours.

## Introduction

Cell migration is involved in many processes such as development, invasion, wound healing, or immune response (Vicente-Manzanares and Horwitz 2011). There is an impressive variety of modalities by which cells migrate, including mesenchymal or amoeboid type of movement for which single cells or a group of cells (Shellard and Mayor 2019) use different propulsive forces for displacement (Othmer 2019). Regardless of the propulsive force or single/collective mode of migration, cells polarize to move (Rappel and Edelstein-Keshet 2017). This is characterized by an asymmetric shape and distribution of proteins, organelles and lipids, as well as differential activities at the two extreme sides of the cell (Vaidžiulyte, Coppey, and Schauer 2019). This polarity allows cells to spatially segregate propulsive and contractile forces in order to move their body forward. In the context of mesenchymal cell migration, the polarity axis of cells is specified by a protruding front and a retracting back (Ridley et al. 2003; Llense and Etienne-Manneville 2015). On the contrary, when cells are not polarized, they present several protruding regions along their contour and barely move (Petrie, Doyle, and Yamada 2009). Several mechanisms have been proposed to explain the long range coordination of front and back activities, from reaction-diffusion of signaling molecules (Jilkine, Marée, and Edelstein-Keshet 2007), cytoskeleton template dynamics (Wang et al. 2013; Gan et al. 2016; Prentice-Mott et al. 2016; Maiuri et al. 2015), mechanical signals such as membrane tension (Houk et al. 2012), to contractility (Schuster et al. 2016; Vicente-Manzanares et al. 2011; Yam et al. 2007; Cramer 2013). Eventually, numerous studies have highlighted the role of retrograde trafficking (Shafaq-Zadah et al. 2016) and directed secretion from the Golgi complex in sustaining persistent migration (Yadav, Puri, and Lindstedt 2009; Yadav and Linstedt 2011; Hao et al. 2020). However, it is not completely understood how these different mechanisms can be combined and what are their respective roles in allowing cells to maintain a stable polarity while migrating.

In the case of mesenchymal migration, cells move thanks to the sum of local protrusive activity (Yamao et al. 2015) and persistent migration relies on lamellipodial persistence (Krause and Gautreau 2014). Protrusions are initiated and controlled by the small RhoGTPases (Jaffe and Hall 2005; Lawson and Ridley 2018). These signaling proteins are engaged in spatiotemporal patterns of activity (Machacek et al. 2009) (Pertz 2010; Fritz and Pertz 2016), thanks to a large set of activators and deactivators, GEFs and GAPs (Bos, Rehmann, and Wittinghofer 2007; Müller et al. 2020). Among the RhoGTPases, Cdc42 has been recognized to be integrated into an excitable signaling network that can spontaneously polarize (Yang, Collins, and Meyer 2015). Cdc42 crosstalks with polarity proteins (Iden and Collard 2008; Etienne-Manneville 2008) and with the cytoskeleton (Bear and Haugh 2014). Notably, persistently migrating mesenchymal cells present a sustained and polarized internal organization, which can be viewed as an ‘internal compass’. This compass corresponds to the polarity axis that can be represented by the axis from the nucleus to the centrosome or the associated Golgi complex (Elric and Etienne-Manneville 2014; Luxton and Gundersen 2011). In wound scratch assay, the Golgi complex reorients in front of the nucleus (Etienne‐Manneville 2006). Similarly, the centrosome reorients toward the leading edge during EMT (Burute et al. 2017). In other studies, the investigators have reported that the Golgi does not align with direction of migration at all (Uetrecht and Bear 2009) or tends to be behind the nucleus when cells are studied on adhesive 1D lines (Pouthas et al. 2008). Thus, the role of the Golgi positioning and the internal compass in persistent migration remains to be clarified.

Based on pioneering work in yeast (reviewed in (Chiou, Balasubramanian, and Lew 2017)), cell polarity could be considered as an emergent property based on the coupling of high-level cellular functions, rather than being attributed to one specific pathway or to one single ‘culprit’ protein (Vaidžiulyte, Coppey, and Schauer 2019). Similarly, the emergence of persistency in cell migration could also rely on the coupling of high-level cellular functions. In the present work, we tested if the coupling between protrusion dynamics and internal cell polarity is present in mesenchymal cells and if this coupling could be sufficient to maintain persistent cell migration. For this, we quantified and manipulated the two subcellular functions described above at short and long timescales. Our experimental results were integrated into a minimal physical model that recapitulates the emergence of persistency from this coupling.

## Results

### Freely migrating RPE1 cells persistently protrude in front of the Golgi

First, we assessed the coupling between the internal polarity axis and cell protruding activity during persistent cell migration. We chose RPE1 cells which are known to have a reproducible internal organization (Schauer et al. 2010) and move persistently (Maiuri et al. 2012). To quantify the orientation of the internal polarity axis of the cell while migrating, we generated stable cell lines, with fluorescently labelled Golgi complex and nucleus. Rab6A fused to a GFP tag was overexpressed to follow the Golgi, and the nucleus was stained with Hoechst 33342 (**see Mat&Meth**). Cell contours were segmented in live by expressing an iRFP-fluorescent reporter anchored to the plasma membrane by a myristoylation motif. The live segmentation was employed to move the stage accordingly to the cell movement in order to keep the cell in the field of view (**sup Figure 1A and Mat&Meth**). This experimental strategy let us image cells with high spatial resolution for up to 16 hrs with a 5-minute temporal resolution (**Figure 1A and Supplementary movie 1**). For each time point, we quantified the direction of movement by measuring the displacement of the center of mass of the cell from the segmented images (orange arrow **Figure 1B**). We quantified the direction of the internal polarity axis by taking the vector joining the centers of mass of the Nucleus to the Golgi (black arrow **Figure 1B**). We then computed the angular difference between the two vectors and averaged it over all time points and over 17 cells. The distribution of the angular difference is sharply pointing toward zero (**Figure 1C**), showing that there is a clear alignment of the Nucleus-Golgi axis with the direction of migration in RPE1 cells.

**Figure 1:**
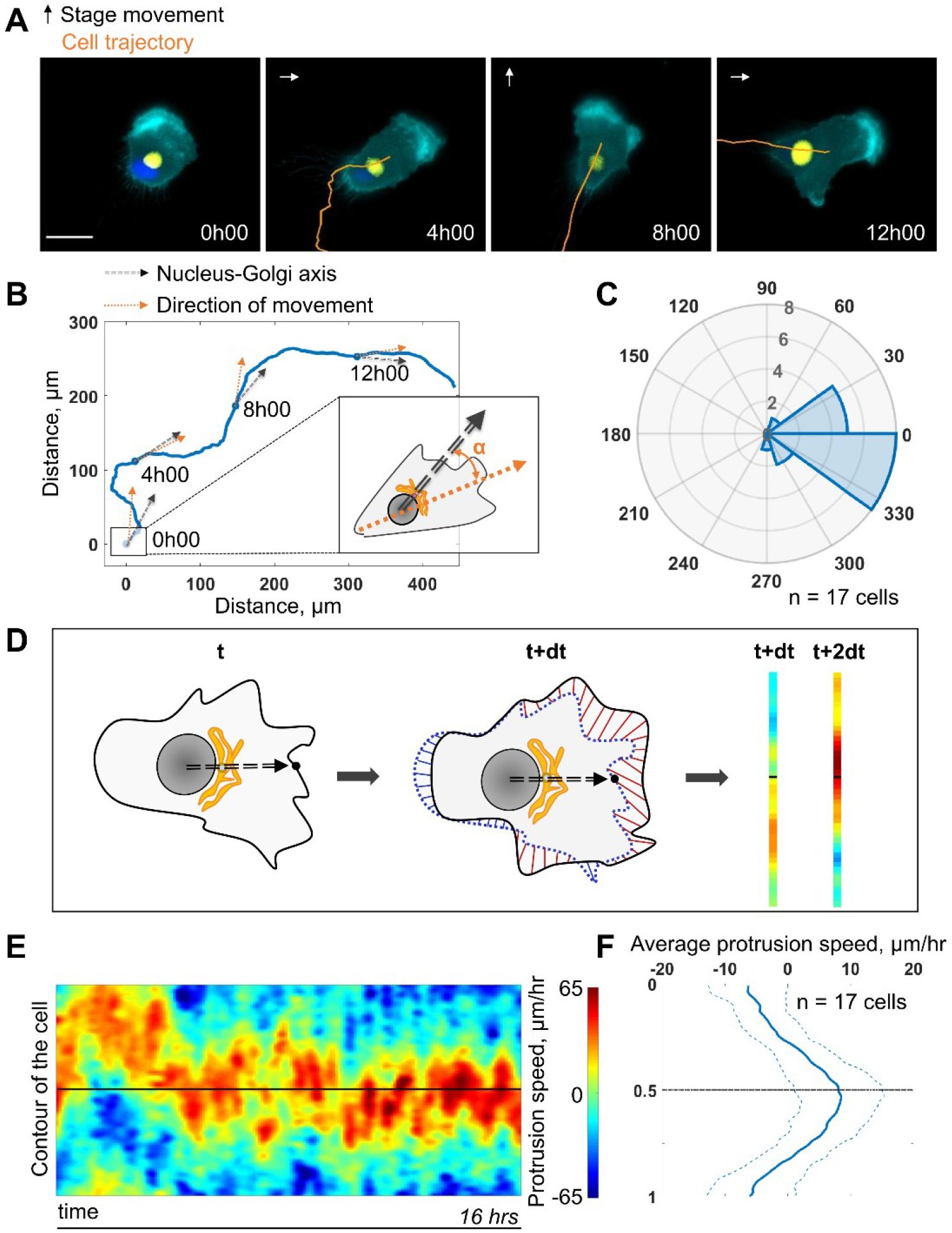
Persistent protrusions form in front of the Golgi complex. **(A)** Snapshots of a representative migrating RPE1 cell at different time points tracked with a feedback routine in which the microscope stage follows a migrating cell (**Supplementary Figure 1A**) for 16 hrs (cyan: myr-iRFP, yellow: GFP-Rab6A, blue: Hoechst 33342, trajectory overlaid in orange, microscope stage movement represented by an arrow, scale bar − 20 μm). **(B)** Full trajectory of a representative cell shown in (a) with Nucleus-Golgi (black dashed arrow) and direction of movement (orange dashed arrow) axes overlaid. **(C)** Polar histogram representing the averaged angle between Nucleus-Golgi axis and direction of movement (n=17 cells). **(D)** Explanatory sketch of how a morphodynamic map of cell shape changes is computed. The contour of the cell is extracted and compared between frames, and stretched out to a line representation, where the distance travelled by a point in the contour is represented (red color meaning protrusion, blue – retraction). **(E)** Morphodynamic map of a representative cell (all maps in **Supplementary Figure 3A**) recentered to Nucleus-Golgi axis (black). X-axis represents time and Y-axis represents cell contour. **(F)** Average protrusion speed over time (n=17 cells, dashed blue line - sd). X-axis represents average protrusion speed and Y- axis represents cell contour with the midline corresponding to **Figure 1E**.

Next, we computed morphodynamic maps of the cell contour, which allows the visualization of protruding activity over time (Machacek et al. 2009). We computed these maps by measuring the displacement of the cell contour between two consecutive time points and by color coding the displacement, from blue (retraction) to red (protrusion) (**Figure 1D, and Mat&Meth**). All displacements along the cell contour (y-axis) were plotted as a function of time. Using the direction of movement as reference (midline on the y-axis) throughout the movement, we found that the protrusive activity is perfectly aligned with the direction of movement (**sup Figure 1C and D**). This showed that cell migration is indeed driven by protrusions. When the Nucleus-Golgi axis is used as reference (midline on the y-axis), the protrusive activity appeared to align well with this axis (**Figure 1E**). Averaging over time and over cells, there is indeed a sustained protrusive activity in front of the Golgi, whose speed is significantly higher than throughout the cell (**Figure 1F**). These results demonstrate that there is a strong correlation between the direction of protruding activity driving cell movement and the orientation of the polarity axis in RPE1 cells indicating a strong coupling of these activities in freely migrating RPE1 cells.

### The Nucleus-Golgi axis does not predict the direction of migration, but aligns when cells move effectively

The correlation we observed does not imply a causal role of the internal polarity axis in driving the persistence of cell migration, because the Nucleus-Golgi axis may follow the direction of migration in a passive manner as a byproduct of cell morphological changes. Thus, we assessed its role by testing if cells start to move in a preferential direction along the given internal polarity axis (**Figure 2A**). For this, we employed the dynamic micropattern technique (Van Dongen et al., 2013) that allows to release cells from a pattern. Cells are initially plated on round adhesive micropatterns coated with fibronectin surrounded by a repulsive PLL-PEG coating. 5 hrs after plating, migration is initiated by adding BCN-RGD that renders the whole surface adhesive. Since the pattern is isotropic, there are no external cues to orient cell escape. We monitored cell movement by tracking the nucleus center of mass, and Nucleus-Golgi axis for 36 cells (**Figure 2B, Supplementary Figure 2A,** and **Supplementary movie 2**). During a first phase of ~5 hrs cells remain on the pattern, and during a second phase they start to move out of it (**Supplementary Figure 2B**). When we compared the orientation of the Nucleus-Golgi axis at t=0 (addition of BCN-RGD) with the direction of escape, we found no correlation (**Figure 2C**). This result shows that the direction of escape is independent from the initial positioning of the Golgi, as previously suggested in the literature (Chen et al. 2013). However, we found a clear correlation between the direction of escape and orientation of the Nucleus-Golgi axis at the time of escape (**Figure 2D**). A detailed temporal analysis (**Supplementary Figure 2C**) showed that both the Nucleus-Golgi axis and cell direction of motion start to align about 2 hours before the escape. Our analysis indicates that they are concomitantly required to initiate effective migration.

**Figure 2:**
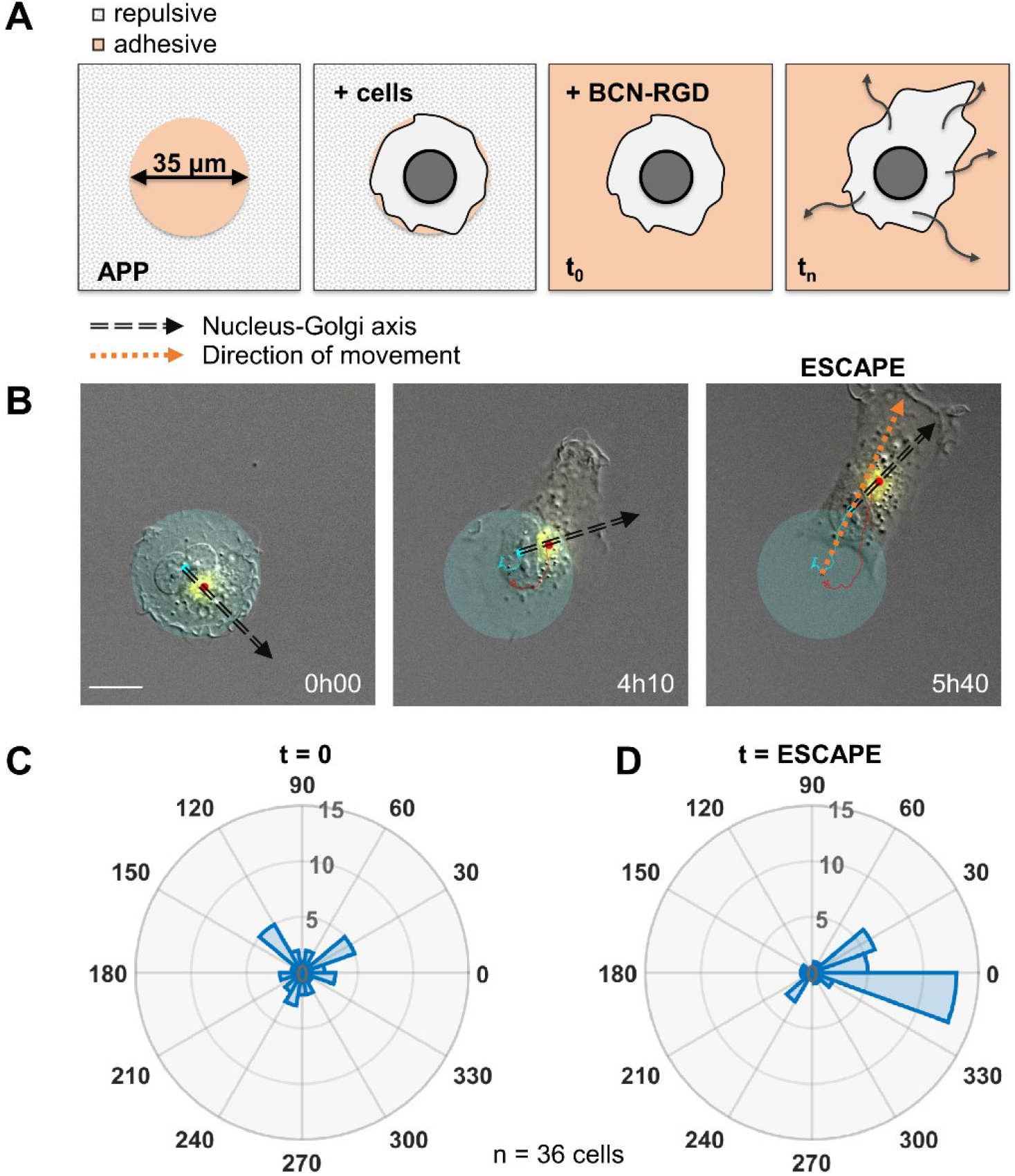
Nucleus-Golgi axis and direction of movement align when a cell starts moving. **(A)** Scheme of the dynamic micropatterns experimental design that is used to study the initiation of cell movement. A cell is confined on a round fibronectin pattern and after the addition of BCN-RGD is enabled to move outside and “escape” the pattern (“escape” is defined to be the moment when the center of the cell nucleus is leaving the area of the pattern). **(B)** Representative RPE1 cell “escaping” the pattern (transparent cyan: pattern, cyan dot: nucleus centroid, yellow: GFP-Rab6A, red dot: Golgi centroid, black dashed line: Nucleus-Golgi axis, orange dashed line: direction of movement, scale bar − 20 μm). **(C-D)** Polar histograms representing the angle between Nucleus-Golgi axis at the beginning of experiment (t=0) (C) or at the time of "escape" (t=ESCAPE) (D) and direction of movement when the cell moves out of the pattern (n=36 cells).

### Disruption of microtubule dynamics abolishes persistence of migration on long timescales

Next, we addressed the role of the internal polarity axis on sustaining protrusion dynamics. Since microtubules (MTs) are known to play a major role in cell internal organization, we perturbed MT dynamics using low doses of Nocodazole (NZ), namely, 0.1 μM, which was sufficient to perturb MT dynamics without impacting cell viability during the experimental set-up (**Supplementary movie 3**). We monitored control and NZ treated cells for 16 hrs using our previously described feedback routine to assess their migration properties. As seen from the corresponding morphodynamic maps, NZ treated cells still show protruding activity that however is less sustained and connected giving rise to separated patches of activity (**Figures 3A-B and Supplementary Figure 3**). We measured the average protrusion speed for each single cell with or without treatment (**Figure 3C**) and found no significant difference, showing that the protrusive ability of NZ treated cells was not altered. As a consequence, the average instantaneous speeds of cells in a short 5 min time window are the same in both conditions (**Figures 3D**). Yet, the directionality ratio -defined as the cell displacement divided by the length of the cell trajectory- strongly differs, pointing to a difference in the directionality of movement (**Figure 3E**). Thus, NZ treated cells are less persistent than control cells, as directly observed from their trajectories (**Figures 3F-G and Supplementary movie 3**). We further quantified the persistence of migration by measuring the autocorrelation of direction of movement, which takes into account only the angle of direction of a moving cell and correlates it over time (Gorelik and Gautreau 2014). The decay of this autocorrelation informs on the timescale over which cells randomize their direction of movement (**Figures 3H**). By fitting an exponential function on the autocorrelation curves of single cells, we extracted a characteristic persistence time of each cell (**Figures 3I**). Control cells are persistent over ~2.5 hrs on average, whereas NZ treated cells are persistent over 20 min on average only. Taken together, our results showed that NZ treated cells are protruding as efficiently as control cells but in a non-coordinated manner over the timescale of more than 20 min. It thus suggests that cell internal organization is required for long-term coordinated protruding activity and persistent cell migration.

**Figure 3:**
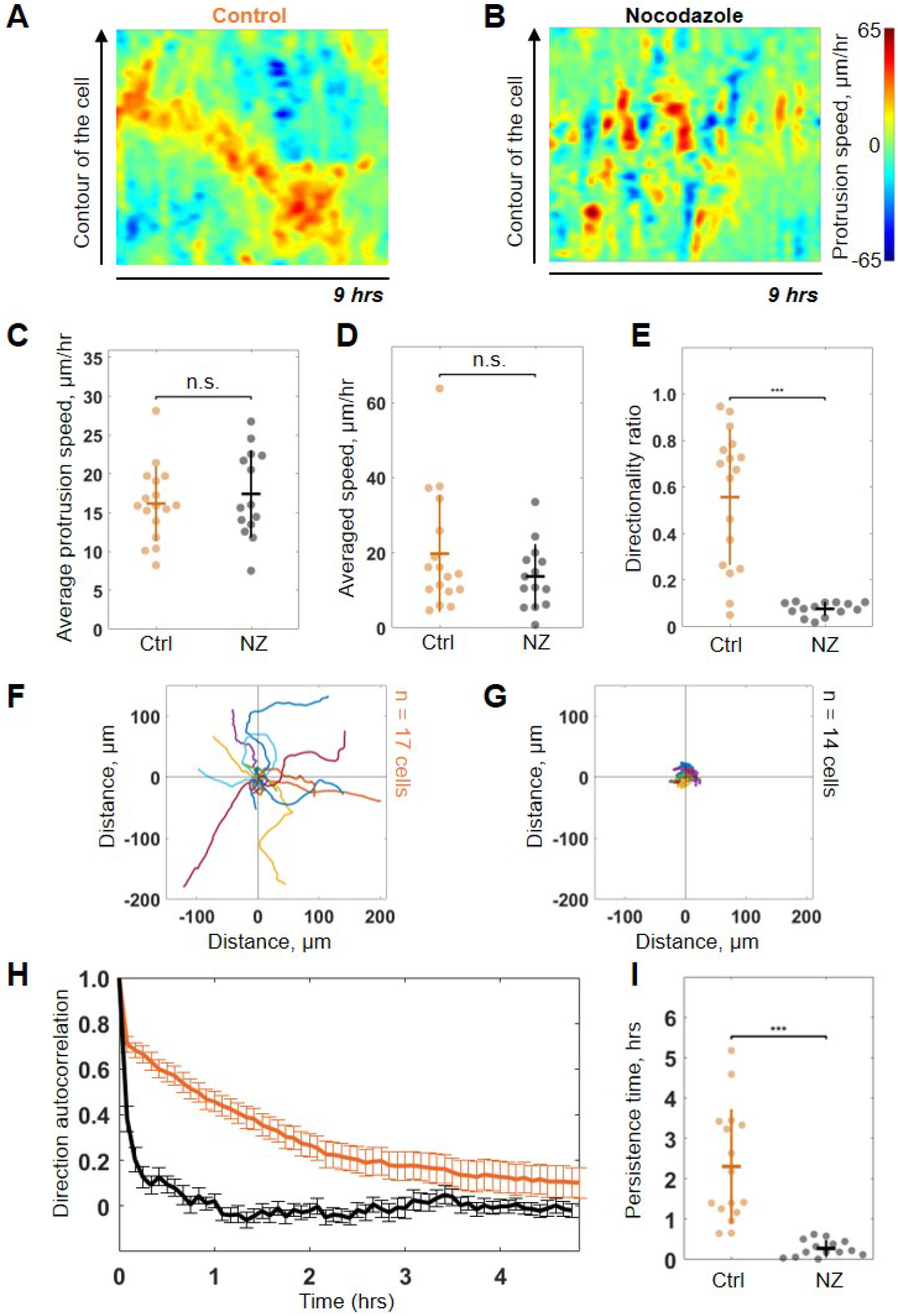
Low dose of Nocodazole reduces persistence of migration. **(A-B)** Representative morphodynamic maps of RPE1 cells freely moving on a fibronectin-covered coverslip in control condition (Ctrl) (A) and with Nocodazole (NZ) (0.1 μm) (B). **(C-E)** average protrusion speed (C), average cell speed (D), and directionality ratio (E) compared in Ctrl and with NZ (Wilcoxon rank sum test, *p ≤ 0.05, **p ≤ 0.01, ***p ≤ 0.001). **(F-G)** Trajectories of RPE1 cells in Ctrl (n=17) (F) and with NZ (n=14) (G) (trajectories plotted over 7 hrs of experiment). **(H-I)** direction autocorrelation (H), and persistence time (I) compared in Ctrl and with NZ (Wilcoxon rank sum test, *p ≤ 0.05, **p ≤ 0.01, ***p ≤ 0.001).

### Persistent cells show sustained polarized trafficking from the Golgi complex to protrusions

The Golgi complex plays important roles in directed secretion of vesicles and cargos to the leading edge of migrating cells (Yadav, Puri, and Lindstedt 2009). To assess the dynamics of Golgi-derived secretion, we followed synchronized secretion of collagen X from the ER to the plasma membrane using the Retention Using Selective Hooks (RUSH) assay. To better visualize the sites of collagen X arrival we combined RUSH with selective protein immobilization (SPI) on the coverslip via antibody capturing that shows secretion of collagen X at 20 min after release from the ER (**see Mat&Meth**) (Fourriere et al. 2019). Interestingly, although we could detect secreted cargos accumulating in the direction of the Nucleus-Golgi axis, (**Supplementary Figure 4A** for representative examples), in 21 of 47 cells, the accumulated cargo did not align with the Nucleus-Golgi axis, probably because we captured cells while turning (**Figure 4A** at 42min when the cargo accumulates in the Golgi). Importantly, we observed the accumulation of secreted cargos toward the newly formed protrusion in these cases (**Figure 4A** at 1hr, **Supplementary Figure 4A** and **Supplementary Movie 4**). To further follow constitutive Golgi-derived trafficking activity at a longer time scale, we used Rab6 as general marker for Golgi-derived secretion (Fourriere et al. 2019). We engineered a CRISPR knock-in cell line with an iRFP fluorescent protein fused to the endogenous Rab6A protein (**see Mat&Meth**). Using HiLo microscopy, we could minimize the signal from the Golgi and enhance signal from the Rab6 vesicles (**Figure 4B, top** and **Supplementary Movie 4**). Moreover, we further suppressed the signal from the Golgi complex by segmenting and masking it to quantify only cell trafficking. We performed the morphodynamic map analysis of cell protrusions (**Figure 4C, top**), in addition to a Rab6- trafficking map (**Figure 4C, bottom**). The latter was computed by measuring the average intensity along lines from the Golgi centroid to the cell contour (**Figure 4B, bottom**). The color code from blue (no Rab6-signal) to red (max Rab6-signal) of this trafficking map shows the hotspots of trafficking as a function of time. We found that the morphodynamic and trafficking maps correlate (**Figure 4D**) confirming a sustained trafficking to protrusions at the long timescale. By performing a temporal cross-correlation analysis, we observed a peak that occurs at a positive time lag of 21±13min indicating that protrusions precede trafficking. This delay is also obvious from the alignment of the morpho and trafficking maps when a cell reorients (e.g. **Figure 4C, black dashed lines**). To further test how internal organization impacts polarized trafficking, we analyzed the secretion of collagen X and the flow of Rab6 positive vesicles in NZ treated cells. As expected, in these cells there was no polarized secretion of vesicles (n=28, see **Supplementary Figure 4B** and **Supplementary Movie 4** for representative examples) and no polarized trafficking of vesicles, but an isotropic directed flow towards the membrane (n=14 cells, see **Supplementary Movie 4** for a representative example). Together, these results show that intracellular polarity axis is required to keep trafficking aligned with protrusive activity over timescales longer than ~20 min.

**Figure 4:**
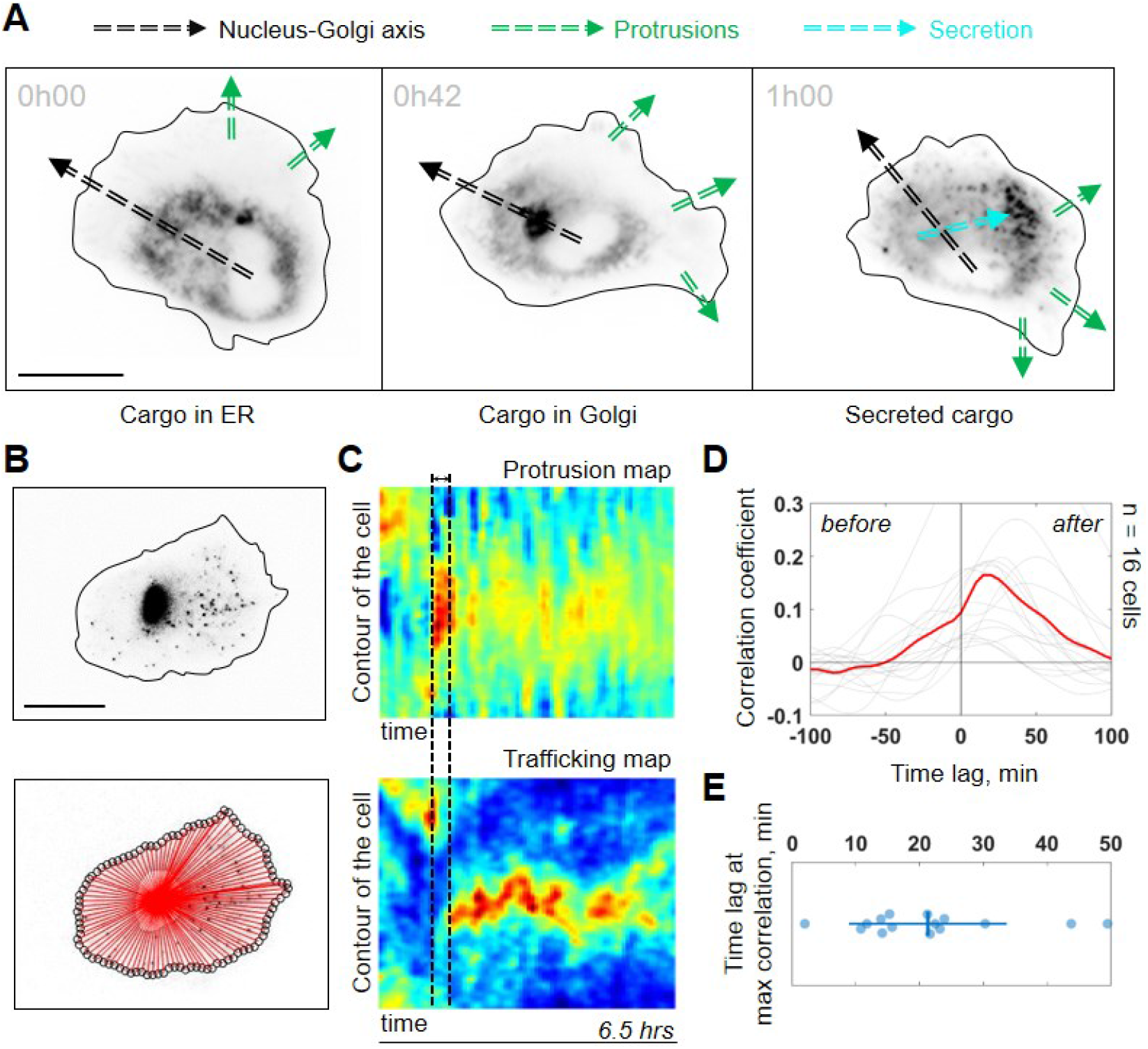
Trafficking from Golgi complex is biased towards the protrusion. **(A)** Collagen X cargo (labelled in black) is travelling from ER to the Golgi complex and secreted during a RUSH assay experiment (green dashed line: protrusions, black dashed line: Nucleus-Golgi axis, cyan dashed line: secretion axis, scale bar - 20 μm). **(B)** Top, RPE1 cells expressing endogenous levels of a marker for post-Golgi vesicles (iRFP-Rab6A) (scale bar - 20 μm). Bottom, red lines represent the lines over which vesicle traffic intensity is calculated over time. **(C)** Top, a representative morphodynamic map representing plasma membrane protrusions in time. Bottom, a representative trafficking map showing the flow of post-Golgi vesicles from the Golgi complex along straight lines towards the surface over time (time period - 6.5 h, black dashed lines represent the time difference between a spike in protrusions (top) and a spike in secretion (bottom). X-axis represents time and Y-axis represents the contour of the cell. **(D)** Cross-correlation coefficient between plasma membrane protrusions and secretion as a function of the time lag (n = 16 cells, red line: average curve depicting the correlation coefficient, grey lines: single cell data; “before” and “after” denote the time before the protrusion peak and after, respectively). **(E)** Single cell time lags in minutes between protrusions and secretion at maximal correlation, obtained by fitting the peak of individual cross-correlation curves (gray curves in (D)).

### An imposed Cdc42 gradient reorients the Golgi complex

Because protrusion activity preceded trafficking from the Golgi, we next investigated how protruding activity regulates internal polarity. We controlled the protrusive activity with optogenetic stimulation while monitoring internal polarity. For the optogenetics, we used the iLID/SspB dimerizing system (Guntas et al. 2015) to locally activate Cdc42 (Valon et al. 2015) by recruiting the catalytic DH-PH domain of ITSN - one of its specific activators- using localized blue light illumination. We used the Nucleus-Golgi axis as a proxy of the internal polarity axis. Using the previously described feedback imaging routine, we added a possibility to induce Cdc42 activation while imaging a migrating cell and adapt the activation pattern to the changing shape of the cell (see **Supplementary Figure 1B** and **Mat&Meth**). Our previous experiments revealed that a sharp gradient of Cdc42 is the most effective to control directionality of cell movement (Beco et al. 2018), therefore, we chose to activate a thin region along the border of the cell. We conducted 16 hrs live-cell imaging experiments, starting with the activation 90°away from the existing Nucleus-Golgi axis. We found that optogenetic activation of Cdc42 was sufficient to reorient the Nucleus-Golgi axis toward the region of activation (**Figure 5A** and **Supplementary Movie 5**). We quantified the rotation of the Nucleus-Golgi axis towards the axis of the optogenetic activation (going from the center of nucleus to the center of activation area) over time for 19 cells and observed a systematic reorientation in 2-4 hours followed by a stabilization of the Nucleus-Golgi axis around 0° (**Figure 5B and Supplementary Figure 5C**). To test the specificity of the Cdc42 activation, we performed control experiments (n=26 cells), in which the DH-PH domain of ITSN is missing (see scheme in **Figure 5C, top**). In control experiments, the Nucleus-Golgi axis constantly moved without stabilization at the axis of optogenetic activation (0°) (**Figure 5C**). To further confirm that the optogenetic activation stabilized the Nucleus-Golgi axis, we optogenetically activated cells in front of existing Nucleus-Golgi axis (**Supplementary Figure 5A**). The axis was stabilized as the angle between Nucleus-Golgi axis and optogenetic activation stayed close to 0° during the full duration of the experiment (n=19 cells) (**Supplementary Figure 5B**). Since our optogenetic activation leads to protrusions and cell migration, the Nucleus-Golgi axis may reorient in a passive manner through cell shape changes. To better control cell shape, we performed similar experiments on round fibronectin micropatterns. Similar to non-patterned cells, the reorientation of Nucleus-Golgi axis aligned with the activation area (**Supplementary Figure 5D-F**). Yet, we found that the Nucleus-Golgi axis reorientation happened faster, with 50% of cells reorienting in 1 hr on a pattern compared to 3 hrs when freely moving (**Supplementary Figure 5C**). Thus, our results show that a biochemical Cdc42 activity but not a change in cell shape is able to reorient the Nucleus-Golgi axis towards it and then stabilize it.

**Figure 5:**
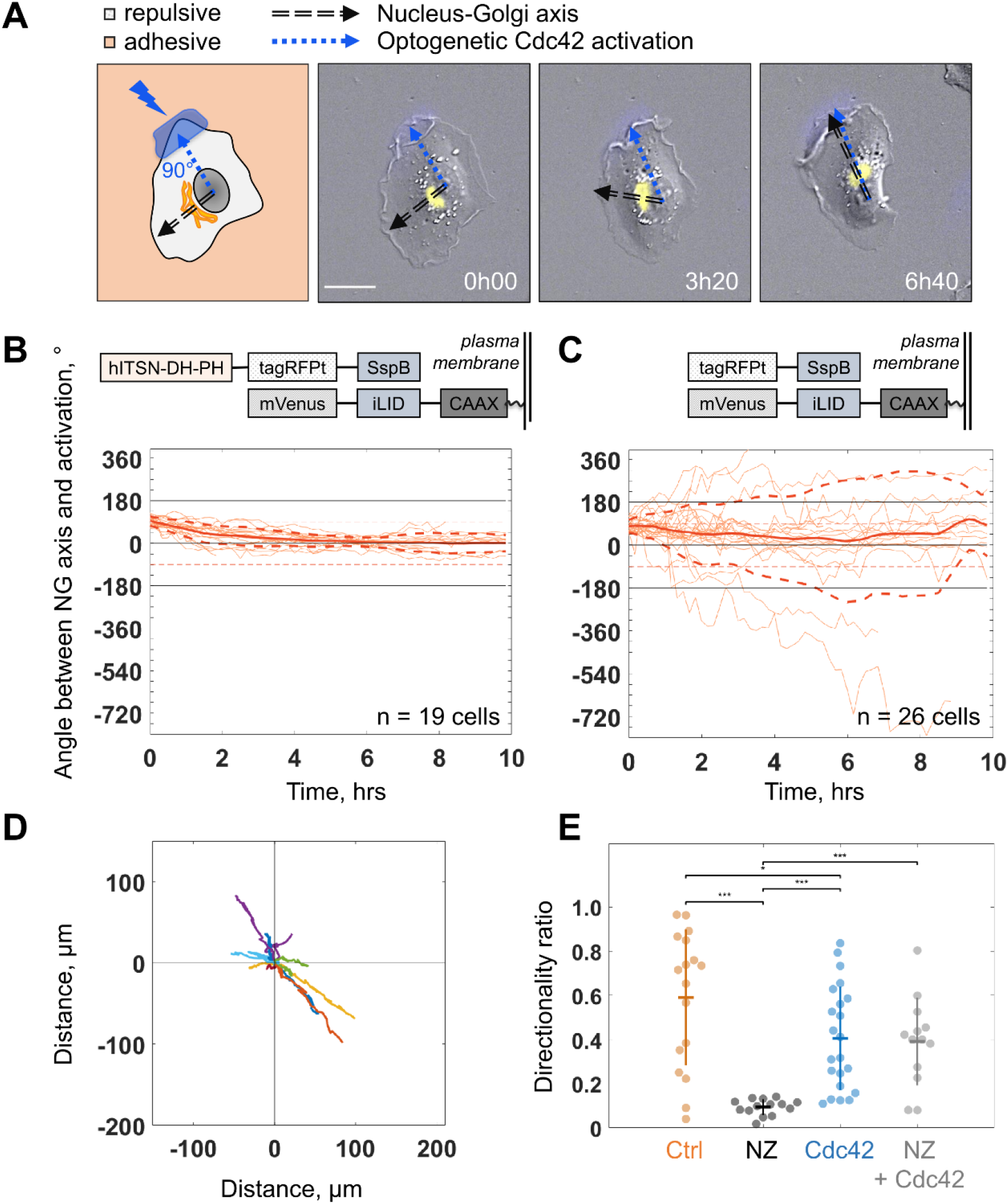
Biochemical gradient of Cdc42 reorients the Golgi complex and rescues directional migration. **(A)** DIC image overlaid with Golgi marker (yellow, iRFP-Rab6A) of an RPE1 cell optogenetically activated 90°away from its initial Nucleus-Golgi axis (black dashed line: Nucleus-Golgi axis, blue dashed line: optogenetic activation axis, scale bar − 20 μm). **(B-C)** Optogenetic activation of Cdc42 90° away from Nucleus-Golgi axis leads to its reorientation in RPE1 cells freely moving on fibronectin covered coverslip (n=19 cells) (B) and is random in control condition (n=26 cells) (C) (thin orange lines: single cell data, thick orange line: data average, dashed thick orange lines: standard deviation; corresponding optogenetic constructs used are depicted above the graphs). **(D)** Trajectories of cells moving in this experimental condition (n=13 cells; trajectories plotted over 7 hrs of experiment). **(E)** Directionality ratio comparison between optogenetically activated cells in presence of Nocodazole (NZ) (orange: freely moving cells (“Ctrl”), black: freely moving cells in presence of NZ (“NZ”), blue: optogenetically activated cells (“Cdc42”), grey: optogenetically activated cells in presence of NZ (“NZ + Cdc42”); Wilcoxon rank sum test, *p ≤ 0.05, **p ≤ 0.01, ***p ≤ 0.001).

### An imposed Cdc42 activation rescues persistent cell migration

We found that persistent migration requires stable protrusive activity, which is lost upon NZ treatment. We thus tested if we could rescue this loss of protrusive stability using activation of Cdc42 by optogenetics. We used optogenetic activation of Cdc42 in presence of NZ (0.1 μM) and found that optogenetic Cdc42 activation indeed rescued the persistence of cell migration, as observed directly from cell trajectories (**Figure 5D**) or from the directionality ratios, which are calculated by taking the ratio of the displacement of the cell and the length of the actual path it took (**Figure 5E**). Whereas directionality ratio in presence of NZ was drastically perturbed, reaching only 19% of directionality in control cells (directionality ratio of 0.11±0.04), optogenetic Cdc42 activation restored the directionality to 0.4±0.2, a number comparable to freely migrating cells (0.59±0.31) and similar to optogenetically activated cells without NZ (0.4±0.24) (**Figure 5E**). The fact that we could rescue persistent migration in NZ treated cells indicates that the loss of persistency in NZ treated cells is the consequence of the absence of a mechanism stabilizing the protrusive activity and not an inherent inability of cells to move persistently.

### A minimal model coupling protrusive activity and polarized trafficking recapitulates persistent migration

Our results indicate that persistent mesenchymal migration emerges from a feedback between the alignment of the internal polarity axis by Cdc42 and stabilization of Cdc42-dependent protruding activity through polarized trafficking towards protrusions. We constructed a minimal physical model to know whether this feedback is enough by itself to recapitulate the features we observed with the persistently migrating REP1 cells (see **Supplementary data** for a detailed explanation of the model). To implement the two sides of the feedback with minimal settings, we chose to model synthetic morphodynamic maps that advantageously capture quantitatively the process of cell migration in a single piece of data.

We first implemented the morphodynamic map corresponding to a single event of protrusive activity, which may comprise several protrusion/retraction cycles. Membrane dynamics following a pulse activation of Ccd42 and Rac1 were previously experimentally obtained and described in (Yamao et al. 2015). In this work, the authors computed the transfer function between a point-like RhoGTPase activity at time 0 and position 0, and the membrane dynamics that follow. We numerically synthetized this transfer function and a Cdc42 pulse of activity extended in space and time (**Figure 6A**) such that the convolution of the two leads to a morphodynamic map similar to a single event of protrusive activity as seen in our data (**Figure 6B**). Next, we simulated full morphodynamic maps by nucleating protrusive events randomly in space and time such that the frequency of protrusive activity of our model matched the data (**Figure 6C**). On these maps, we assumed that the cell possessed an internal polarity axis parametrized by a moving point, *c_axis_*(*t*), on the y-axis, which corresponds to the intersection between the polarity axis and the cell contour. For sake of simplicity, we did not make the distinction between the axis of directed trafficking/secretion and Nucleus-Golgi axis but considered a single effective one. We then implemented the feedback between polarity axis and protrusion dynamics. For the first side of the feedback, we introduced a probability *P_polarized_* to nucleate a protrusive event in front of the polarity axis and a probability *P_rand_* = 1 − *P_polarized_* elsewhere. When *P_polarized_* = 0, protrusions are happening randomly along the contour, and when *P_polarized_* = 1, protrusions are always happening in front of the polarity axis (**Figure 6D**). For the second side of the feedback, we assumed that the polarity axis was pulled towards the protrusion by an effective force *F_prot_* that acts against a force *F_basal_* characterizing the random rotation of the polarity axis. The bias of the protrusive activity on the polarity axis positioning can be parametrized by a number *κ* such that 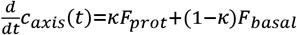. When *κ* = 0, the polarity axis follows its natural evolution, and when *κ* = 1, the polarity axis follows the protrusive activity (**Figure 6E**). The strength of both sides of the feedback can thus be summarized by two numbers between 0 and 1. For a given value of these two numbers, we could simulate realistic morphodynamic maps ranging from non-persistent to persistent migrating cells (**Figure 6F** and **Supplementary movie 6**). From these maps, we generated cell trajectories from which we computed the autocorrelation of direction and persistence time, as for our experimental data (**Figure 6F**). In addition, we computed two other independent parameters aimed at quantifying cell polarity (**Figure 6G**). The protrusive unicity index characterizes how many distinct protrusive activities are competing at a given time, and the alignment index characterizes how well the polarity axis aligns with the direction of movement.

**Figure 6:**
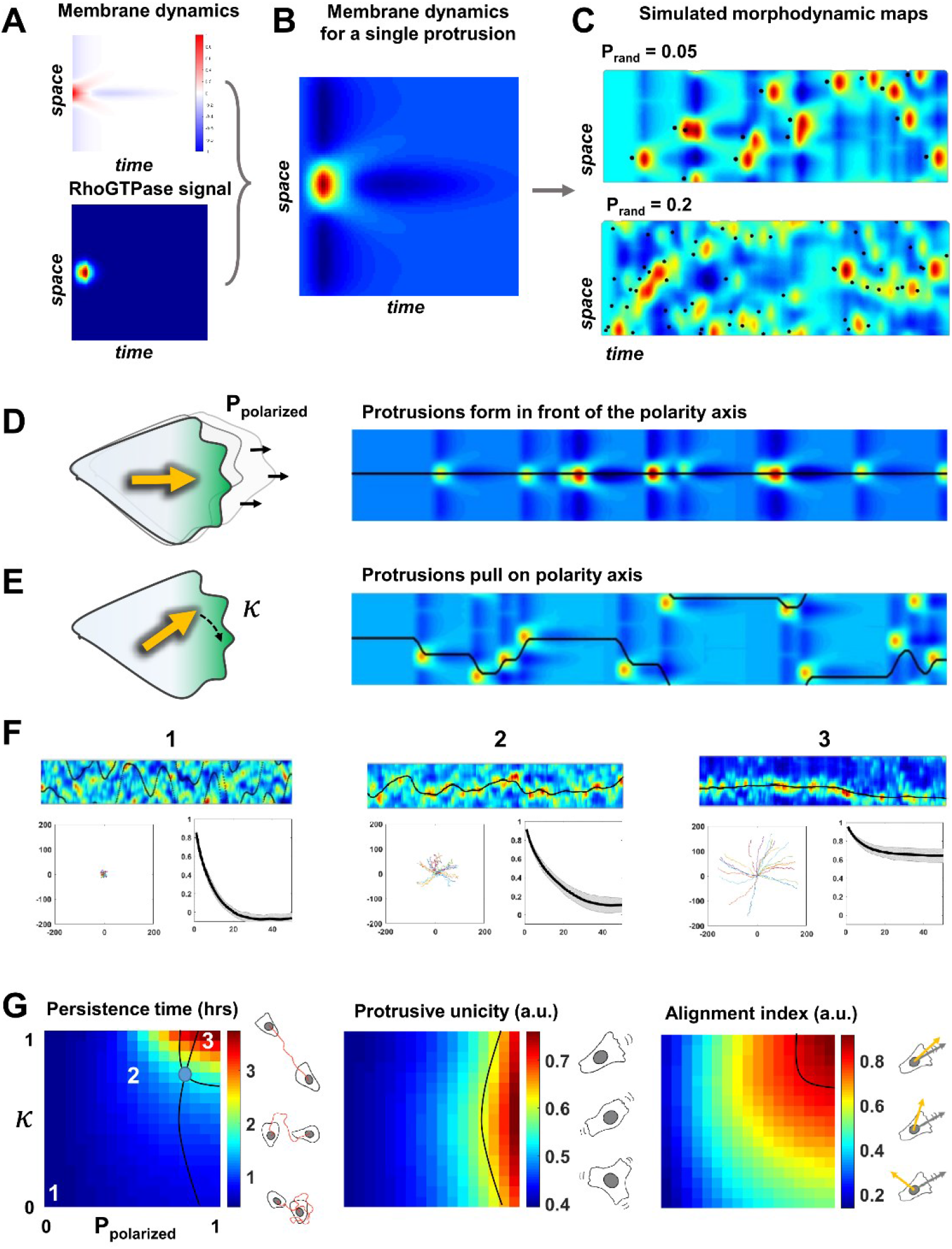
A minimal physical model based on the coupling between protrusive activity and internal polarity recapitulates persistent migration. **(A-B)** Membrane dynamics for a point-like Cdc42 activation (A, upper panel) were convolved with a RhoGTPases signal (A, lower panel) to compute membrane dynamics for a single protrusive event (B). **(C)** Overall synthetic morphodynamic maps generated by varying the protrusive activity frequency. The intensity of protrusive activity varies from one protrusion every 20 frames (top), to one every 5 frames (bottom), the latter value having been retained for our simulations in (F-G). **(D-E)** Implementation of the feedback, with *P_polarized_* quantifying the probability to form a protrusion in front of the polarity axis (equal to 1 in (D)), and *κ* quantifying the capacity of protrusions to pull on the polarity axis (equal to 1 in (E)). **(f)** Examples of morphodynamic maps (top, black line is *c_axis_*(*t*) the direction of the internal polarity axis), cell trajectories (bottom left), and autocorrelation of direction (bottom right) for different values of strength of the feedback. 1: *P_polarized_* = 0; *κ* = 0.2: *P_polarized_* = 0.5; *κ* = 0.8. 3: *P_polarized_* = 0.8; *κ* = 1. **(G)** Phase diagrams of persistence time, protrusive unicity, and alignment index. Black lines in the second and third diagrams correspond to the experimental values (protrusive unicity ~0.6 and alignment index ~0.7). These lines are reported on the first diagram, where they cross at a value of persistence time (blue dot) of ~2 hrs, consistent with our experimental measurement.

Running our simulation for all possible values of *P_polarized_* and *κ*, we obtained three phase diagrams for the persistence time, protrusive unicity, and alignment index (**Figure 6G**). These ‘look-up’ tables differ in their dependencies with regards to the two parameters, and can, thus, be used to estimate their values independently. When combined, they should converge to a single couple of values. If it is the case, it would be a signature of the consistency of our minimal model. It was indeed the case for our data on RPE1 cells, where we found a persistence time of 2.3±1.4 h, a protrusion unicity index of 0.63±0.12, and an alignment index of 0.72±0.21. Using these numbers and the phase diagrams to estimate *P_polarized_* and *κ*, we obtained a region of the parameter space that is consistent and predicts that *κ* = 0.9 and *P_polarized_* = 0.7. Thus, our model suggests that the high persistence of RPE1 cells can be explained by a relatively high values of the feedback strengths. The effect of NZ can be captured by assuming that *P_polarized_* = 0, such that cells become persistent only over the time scale of a single protrusive activity (20 min). Interestingly, RPE1 cells sit in the persistence time phase diagram at the relatively sharp transition between non-persistency and super-persistency. This suggests that cells might be tuned at an optimal functioning point, to be persistent in their migration while not being locked into a straight path, possibly to be able to respond to environmental cues.

## Discussion

In this work, we propose that persistent mesenchymal migration observed on a timescale of several hours can emerge from a feedback between protrusion dynamics and polarized trafficking. This feedback mechanism corresponds to the one that has been intensively documented in yeast, where polarized bud formation is dynamically maintained by coupling of transport and signaling (Eugenio et al. 2008). Using experimental approaches, we first showed that protrusion dynamics and polarized trafficking are coupled in mesenchymal RPE1 cells. We demonstrated using optogenetics that sustained local activation of Cdc42 is sufficient to reorient the Nucleus-Golgi axis in 2-4 hours (**Figure 5**). Moreover, we showed that the Nucleus-Golgi axis correlated with the direction of migration and the trafficking of Rab6 secretory vesicles in freely migrating cells (**Figure 1 & 4**). Our optogenetic result thus suggests that a sustained protrusive activity does reorient the trafficking and secretory pathway towards protrusions. We observed a time lag of 20 min between protrusions and redirection of the trafficking of Rab6-positive vesicles and that secretion was preferentially directed to newly formed protrusions. Taken together, these results strongly support the fact that protrusions orient polarized trafficking on a short time scale, and orient the Nucleus-Golgi axis on a longer one.

On the other hand, our data demonstrate that polarized trafficking sustained protruding activity: using low doses of Nocodazole to disrupt internal cell organization and polarized trafficking (**Supplementary Figure 4B** and **Supplementary Movie 4**), we showed that the persistence time of cell migration dropped from 2.3 hours to 20 minutes (**Figure 3**). Interestingly, protrusion speed and instantaneous cell speed were not affected, showing that the loss of persistency was not due to cell’s inability to protrude or move. Our rescue experiment of constant Cdc42 activation (**Figure 5**) confirmed this fact. Of note, NZ treated cells were still persistent over 20 minutes, showing that in this condition the protrusive activity is stable for a longer time than the duration of a single protrusion-retraction event (on the order of 100 s). Thus, NZ treated cells are still able to stabilize a protrusive activity over several cycles, possibly thanks to the existence of a vimentin template (Gan et al. 2016). However, NZ treated cells were not able to stabilize their protrusive activity over longer time, which we attributed to the loss of polarized trafficking (**Supplementary Movie 4**), potentially of protrusion-promoting factors toward the cell front in Rab6-positive vesicles (**Figure 4**). Rab6 has been proposed to be a general regulator of post-Golgi secretion and it has been shown that irrespective of the transported cargos most Rab6-positive carriers are not secreted randomly at the cell surface, but on localized hotspots juxtaposed to focal adhesions (Fourriere et al. 2019). However, we cannot exclude that protrusion-promoting factors are transported from other compartments, such as the recycling endosomal compartment that is found at the proximity of the Golgi complex.

We recapitulated the two sides of the feedback in the framework of a new minimal physical model. This model is based on the coupling between an internal polarity axis - a vector, and protrusion dynamics modelled by synthetic morphodynamic maps. This model can be thought of as a ‘cell compass’, where protrusions would pull on the needle that has some inertia, and the direction of the needle would locally promote the initiation of protrusions. Many mathematical models of cell polarization (Mogilner, Allard, and Wollman 2012; Jilkine and Edelstein-Keshet 2011) or of cell migration (Danuser, Allard, and Mogilner 2013) were previously introduced, but our approach differs in the sense that the whole cell migratory behavior can be described by only two effective parameters that quantify the strength of the feedback. We showed that the persistence time of RPE1 cells can be predicted from the measurement of two parameters - the average number of competing protrusions and the alignment of polarity axis with direction of motion (**Figure 6**). It would be interesting to see whether other cell types can also be consistently placed in our phase diagrams. Our model shows that cells can polarize with a unique protruding front (protrusive unicity close to 1) when the probability to form protrusion in front of the polarity axis is high (P_polarized = 1). Of course, this was expected since we assumed that there was a unique polarity axis. In our experiments, we indeed observed a unique axis of polarized trafficking (**Figure 4**), and it remains to be understood how cells achieve this unicity. As a matter of fact, we could imagine that multiple protrusions would be sustained simultaneously, associated to their own polarized trafficking routes, and with the same feedback mechanism being involved. A possible answer would be the existence of a limiting component in the system, as proposed in the context of yeast polarity (Chiou, Balasubramanian, and Lew 2017). Alternatively, the level of RhoGTPase activity might be tuned to limit the number of competing protrusions, as suggested by the relationship between Rac1 activity and directional persistent migration (Pankov et al. 2005).

To conclude, our present work focused on the coupling between protrusion dynamics and polarized trafficking. Many other functional units supporting cell polarity are likely to be involved (Vaidžiulyte, Coppey, and Schauer 2019), and it will be interesting to see in future studies how other coupling mechanisms can contribute to the robustness of cell polarity during persistent migration. Additionally, it would be of interest to see how our conclusions can be extended to other types of migration, such as the amoeboid one.

## Supporting information

Supplementary Material with movie legend

Supplementary Material - Model

## Acknowledgements

We thank the Institut Curie Cytometry platform for cell sorting; Remy Fert and Eric Nicolau from the Mechanical Workshop for their technical assistance; Gaëlle Boncompain for sharing the RUSH system contructs; Bruno Goud for critical reading of the manuscript and helpful discussions; John Manzi and Fahima Di Federico from UMR168 BMBC platform, Maud Bongaerts and Laurence Vaslin for experimental advice and help with plasmid constructs; Jean de Seze for fruitful discussions and help with adaptation of the cell tracking routine. We acknowledge financial support from the Labex CelTisPhyBio (ANR-10-LBX-0038), the Labex and Equipex IPGG (reference: ANR10-NANO0207), and Idex Paris Sciences et Lettres (ANR-10-IDEX-0001-02 PSL), as well as the Centre National de la Recherche Scientifique and Institut Curie. French National Research Infrastructure France-BioImaging (ANR10-INBS-04). Institut Convergences Q-life (ANR-17-CONV-0005). K.V. was supported by Programme doctoral Interface pour le Vivant (IPV) and Fondation pour la Recherche Médicale (FRM) (FDT201904008167).

## Author Contributions

Conceptualization, K.V., K.S. and M.C.; Methodology, K.V., K.S. and M.C.; Software, K.V., A-S.M., W.B. and M.C.; Formal analysis, K.V. and M.C.; Investigation, K.V.; Modelling, M.C.; Resources, A.B.; Writing—original draft, K.V. and M.C.; Writing—review and editing, K.V., A-S.M., K.S. and M.C.; Visualization, K.V., A-S.M. and M.C.; Supervision, K.S. and M.C.; Funding acquisition, K.V., K.S. and M.C.

## Materials and Methods

### Cell culture

hTERT RPE1 cells (CRL-4000 strain, ATCC, Manassas, VA) were cultured at 37°C with 5% CO_2_ in Dulbecco’s modified Eagle’s/F-12 medium supplemented with 10% fetal bovine serum, GlutaMAX (2 mM) and penicillin (100 U/mL)-streptomycin (0.1 mg/mL). Cells were passaged twice a week in a ratio of 1/10 by washing them with PBS (1x) solution and dissociating using TrypLE Express (Thermo Fisher Scientific, Waltham, MA) reagent for 5 min.

### Plasmids, transfection, and stable cell lines

#### Plasmids

pLL7.0: hITSN1(1159-1509)-tgRFPt-SSPB WT (Plasmid #60419), pLL7.0: Venus-iLID-CAAX (from KRas4B) (Plasmid #60411), pMD2.G (Plasmid #12259), psPAX2 (Plasmid #12260) lentiviral plasmids were bought from Addgene (Watertown, MA). pHR: myr-iRFP plasmid was a gift from Simon de Beco (Institut Curie, France) and pIRESneo3: Str-KDEL-ST-SBP-EGFP (Plasmid #65264) was a gift from Gaëlle Boncompain (Institut Curie, France).

#### Transfection

Transfections were performed using X-tremeGENE 9 (Roche Applied Science, Penzburg, Bavaria, Germany) according to the manufacturer’s instructions using an equal amount of plasmid DNA for each construct (1 μg) and a ratio of 3:1 of transfection reagent and DNA.

#### Stable cell lines

Stable cell lines were generated using two techniques - lentiviral infection and CRISPR cell line development.

##### Lentiviral

For lentivirus production, packaging cell line HEK 293T cells were cotransfected with pMD2g (envelope), psPAX2 (packaging) and lentiviral (transfer*) plasmids in a 1:3:4 ratio, respectively (*pHR-, pLVX- or pLL7-based plasmids were used as transfer plasmids). Lentivirus was harvested 48 h after transfection and filtered from the supernatant of cell culture by passing it through 0.45 μm filter using a syringe. Next, the target RPE1 cell line was transduced for 24 h with media containing lentiviral particles. Subsequently, RPE1 cells were selected by Fluorescence-activated Cell Sorting (FACS) according to the fluorescence level of transduced protein.

##### CRISPR

A CRISPR approach was used to develop a cell line with a heterozygous iRFP-Rab6A knock-in as a Golgi complex label. CRISPR sgRNAs were designed using the Optimized CRISPR Design tool CRISPOR TEFOR (for sequences see the table below). For sgRNA-encoding plasmids, single-stranded oligonucleotides (Eurofins Genomics, Germany) containing the guide sequence of the sgRNAs were annealed, phosphorylated and ligated into BbsI site in px335 plasmid, coding for Cas9 (kindly provided by M. Wassef, Curie Institute, Paris, France). Homology arms of ~800 bp were amplified from genomic DNA using PCR primers with 40 bp overhangs compatible with pUC19 backbone digested with Xba1 and Ecor1 (New England Biolabs, Ipswich, MA) (sequences in the table below). Gibson reactions were performed using a standard protocol with home-made enzyme mix (Gibson, 2009). RPE1 cells were transfected with 90 μL of Polyethylenimine (PEI MAX #24765 Polysciences, Warrington, PA) and 15 μg of the pX335-gRNA and pUC19-homology arms-iRFP plasmids, both diluted in 240 μL NaCl 150mM. 7 days after transfection, positive cells were sorted with FACS for enrichment, and after additional 10 days, FACS sorted again by single cell per well in a 96-well plate. All 96 clones were screened by PCR and 8 clones were selected for further verification by Western blotting, followed by sequencing.

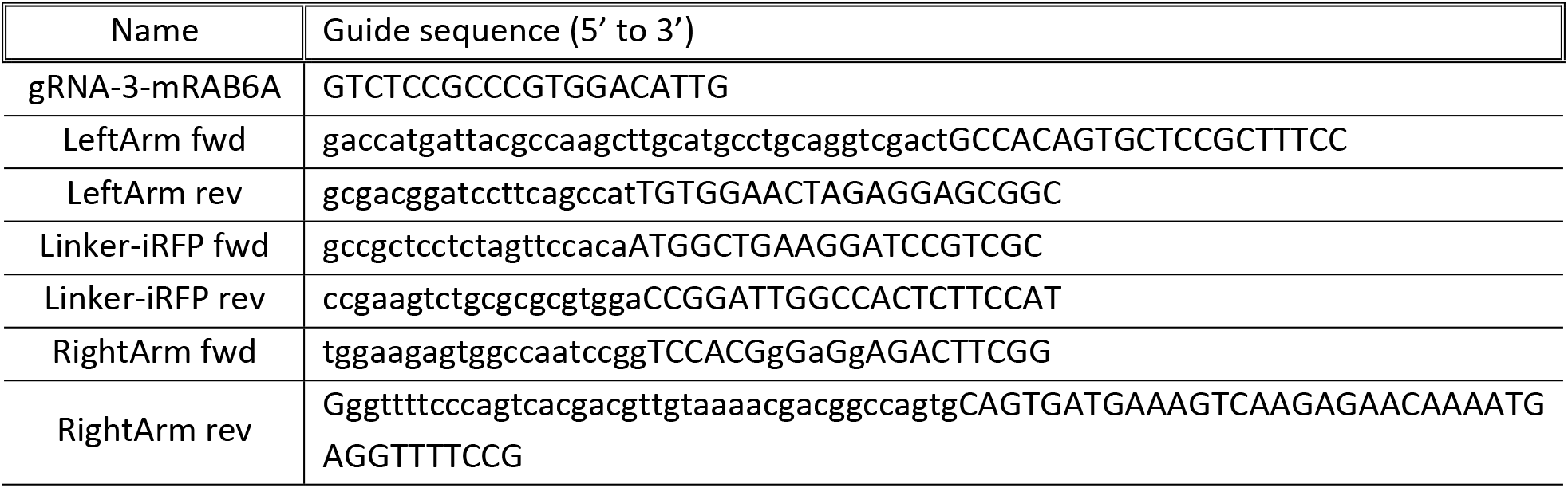

### Micropatterning

#### Coverslip preparation

Coverslips for live-cell imaging were prepared by cleaning round glass coverslips (d=25 mm, 0.17 mm thickness) (Menzel Gläser, Thermo Fisher Scientific, Waltham, MA) for 1 min in O_2_ plasma and incubating them with fibronectin (2 μg/mL) (Merck, Darmstadt, Germany) in 100 mM NaHCO_3_ (pH 8.5) for 1 h in room temperature. Coverslips were washed with PBS (1x) three times and stored in + 4°C in PBS (1x).

#### Static pattern

Micropatterned coverslips were prepared as described by (Azioune et al. 2009): O_2_ plasma-cleaned coverslips were incubated with 100 μg/mL of PLL-g-PEG (Surface Solutions, Switzerland) in 10 mM HEPES, pH 7.4 for 1 h. They were then exposed to deep UV through micropatterned quartz/chrome photomasks (Toppan, Round Rock, TX) for 5 min, and incubated with fibronectin (20 μg/mL) in 100 mM NaHCO_3_ (pH 8.5) for 1 h.

#### Releasable (dynamic) patterns

Releasable micropatterns were prepared similarly, with PLL-g-PEG being replaced by azido-PLL-g-PEG (APP) at 100 μg/mL and fibronectin used at lower concentration (10 μg/mL). Migration was released by addition of 20 μM BCN-RGD for 10 min (described in (van Dongen et al. 2013)).

### Drug assays

In drug assays, RPE1 cells were treated with Nocodazole (0.1 μM) (Sigma-Aldrich, St.Louis, MO) and Golgicide A (35 μM) (Sigma-Aldrich, St. Louis, MO) in complete DMEM/F-12 medium at 37 °C in 5% CO_2_ for the duration of the experiment. 30 min before the experiment and before addition of the drug, cells were incubated with Hoechst 33342 (1 μg/ml) dye (Thermo Fisher Scientific, Waltham, MA), to label cell nuclei. Then, the dye was washed with 1X PBS buffer (pH = 7.5) and a drug, diluted in complete medium, was added. Cells were imaged immediately after addition of a drug.

### RUSH and SPI assays

#### RUSH assay

Retention using selective hooks (RUSH) assay was performed as described in (Boncompain et al. 2012) and (Boncompain and Perez 2012). RPE1 cells were transfected with 2 μg of plasmid containing the RUSH system (Str-KDEL-SBP-EGFP-Col10A1) and a GFP-labelled Collagen type X (ColX) cargo. 24 h after transfection, cells were put on anti-GFP antibody coated glass coverslips and let to attach. After 2 h, biotin was added (40 μM final concentration from 4 mM stock) to the full medium, triggering the release of the cargo. Cells were imaged for 2 h, until the ColX cargo passed from the endoplasmic reticulum (ER) to the Golgi complex and then was secreted. Coverslip being covered with anti-GFP antibodies enabled the GFP-labelled ColX cargo capture upon secretion.

#### SPI assay

Selective protein immobilization (SPI) assay was performed as described in (Fourriere et al. 2019). Round glass coverslips (d = 25 mm) were either autoclaved and incubated in bicarbonate buffer (0.1 M pH 9.5) for 1 h at 37°C (300 μL, upside down) or plasma cleaned (2 min vacuum, 1 min plasma). Next, the coverslips were transferred to poly-L-lysine (0.01% diluted in water) and incubated for 1 h at 37°C (300 μL, upside down). After being washed in 1X PBS and dried, they were transferred to a solution of anti-GFP antibodies (diluted in bicarbonate buffer) and incubated for 3 h at 37°C (70 μL, upside down). After another wash with PBS, cells were seeded on top of coated coverslips in complete medium (at least 2 h given for cells to attach before conducting a RUSH assay). Antibodies used for coating in this study were rabbit anti-GFP (A-P-R#06; Recombinant Antibody Platform of the Institut Curie; dilution 1:100).

### Optogenetics

For local subcellular activation of a RhoGTPase Cdc42, an optogenetic dimer iLID-SspB was used as described in (Guntas et al. 2015). It was activated by illumination with blue light (440±10 nm). RPE1 cells used in the optogenetic experiments were engineered to stably express the optogenetic dimer and selected for average-high fluorescence level by FACS (the highest expressing cells were discarded, as they were not responsive to optogenetic activation). Experiments were performed in live-cell imaging conditions described in the paragraph “Imaging”, using a DMD projector and a blue (440±10 nm) LED illumination source. The projection of blue light was controlled with an interface of a MATLAB script and a microscope controlling MetaMorph software by sending a static pattern of light, or using the imaging routine described below. The illumination pattern was optimised for a local signal and weak illumination to reduce the phototoxicity and enable long-term experiments.

### Imaging

#### Live-cell imaging

All imaging was performed at 37° C in 5% CO_2_ with an IX71 inverted fluorescence and Differential Interference Contrast (DIC) microscope (Olympus, Melville, NY), controlled with MetaMorph software (Molecular Devices, Eugene, OR). The microscope was equipped with a 60x objective (NA=1.45), motorized stage and filter wheel with SmartShutter Lambda 10-3 control system (Sutter Instrument Company, Novato, CA), a stage-top incubation chamber with temperature and CO_2_ control (Pecon, Meyer Instruments, Houston, TX), ORCA-Flash4.0 V3 Digital CMOS camera (Hamamatsu Photonics K.K., Japan), z-axis guiding piezo motor (PI, Karlsruhe, Germany), CRISP autofocus system (ASI, Eugene, OR), a laser control system with azimuthal TIRF configuration (iLas2, Roper Scientific, Tucson, AZ) and a DMD pattern projection device (DLP Light Crafter, Texas instruments, Dalas, TX), illuminated with a SPECTRA Light Engine (Lumencor, Beaverton, OR) at 440±10 nm. Before imaging, cells were dissociated using Versene Solution (Thermo Fisher Scientific, Waltham, MA) and seeded for adhesion on previously mentioned prepared coverslips.

#### TIRF microscopy

Total Internal Reflection Microscopy (TIRF) was used to excite a thin band of fluorophores close to the membrane of adherent cells and avoid out-of-focus fluorescence (Mattheyses, Simon, and Rappoport 2010). A variation of TIRF, called azimuthal TIRF, was used to generate homogeneous illumination and to avoid fringe interferences and imaging artefacts.

#### HILO microscopy

Highly inclined and laminated optical sheet (HILO) microscopy was used to illuminate the cell at an angle with a thin inclined beam, which increased the signal-to-noise ratio (SNR) when imaging the Golgi complex and nuclear markers in the cell.

### Feedback routine and DMD illumination

#### Feedback routine

Imaging feedback routine to follow migrating cells with high magnification (60x) was established by using a combination of scripts in MetaMorph and MATLAB. It ensures that the microscope stage moves together with a moving cell, always keeping it in the field of view. The main script was written in MATLAB, which commands MetaMorph through calling its macros called ‘journals’. It enables imaging of multiple stage positions (i.e. multiple cells) in multiple wavelengths in one experiment. It can be controlled with a GUI, which displays a selected position and its coordinates and pattern of activation for every cell. The amount of acquisition channels and timing can be selected globally for the full set of cells in the experiment. One specific wavelength is chosen as segmentation channel. The images from this channel are used to segment the shape of the cell and to instruct its position. The segmentation threshold can be adjusted for every position and the watershed algorithms can be chosen to separate two touching objects (i.e. cells) in every case.

#### DMD illumination

Local subcellular activation with light for optogenetic experiments was achieved by a Digital Micromirror Device (DMD) (Davis 2013) with dimensions of 640 × 480, able to generate 8-bit grayscale patterns. The pattern was individually adjusted for every cell and dynamically evolved during the experiment according to the cell shape. The activation step was incorporated in the previously described imaging feedback routine.

### Image analysis

#### Image segmentation

Live-cell imaging data obtained using our cell tracking feedback routine was analyzed using a custom-built Matlab script, which allowed segmentation of the shape of the cell, the Golgi complex and the nucleus, tracking of their position and visualization of their trajectories. All three structures of interest (cell membrane, the Golgi complex and the nucleus) were fluorescently labelled, so the shape segmentation was done by using a Matlab function ‘graythresh’ to select the pixels over a certain threshold of fluorescence intensity. Next, the image was binarized with a function ‘im2bw’, structures touching the image border were deleted with ‘imclearborder’, small objects were removed from the image with ‘bwareaopen’, stuctures were closed by dilate-erode with ‘imclose’, using a disk with a radius of 10 pixels as a structuring element, and the holes in the structure were filled with ‘imfill’ function. The resulting segmented cell shape was then used to extract the ‘centroid’ of the cell by ‘regionprops’ function. The same procedure was used to segment the Golgi complex and the nucleus shapes, but with slightly different threshold range. In addition to that, the centroid search area was optimized by selecting the centroid closest to the centroid in the previous image, which is particularly useful when several regions of interest are found in the image. The extracted centroids of all three structures were then concatenated into trajectories depicting the movement through the full experiment. Using our tracking routine implies that the microscope stage would move when a cell moves out of the defined field of view. In this case, the trajectory presents jumps, due to the stage movement. These jumps were corrected using the recorded positions of the stage and a defined scaling parameter. The corrected real trajectories were then used for further analysis.

#### Nucleus tracking (semi-manual)

Excitation with blue light had to be avoided in optogenetic experiments because of optogenetic system’s sensitivity to blue light, so the nuclei in these experiments were tracked semi-manually in the DIC channel, using another custom-made Matlab routine. The estimated center of the nucleus was manually chosen by single-clicking on the image, the centroids of the nuclei were recorded, concatenated into a trajectory and corrected according to the stage movement.

#### Reorientation plot analysis

Evaluation of the Nucleus-Golgi axis reorientation requires two axes: one going through the centroid of the Nucleus and the centroid of the Golgi and another one going through the center of the Nucleus and the center of the optogenetic activation area. The previously described image segmentation techniques were used to segment the optogenetic activation area. Next, the angle between the two axes was calculated using centroid coordinates and an inverse tangent function ‘atan’ in Matlab. Then, the angle in radians was wrapped to [0 2pi], using the ‘wrapTo2Pi’ function and unwrapped with ‘unwrap’ function. The angle, then, was converted from radians to degrees and plotted in a graph.

#### Cumulative plot analysis

The speed of the axis reorientation process in different experimental conditions was compared by plotting the data from the previously described reorientation analysis. The first time point, when the angle reached 30° was chosen to delineate that the Nucleus-Golgi axis has reoriented, which gave the timing of the reorientation for each cell. Next, the ‘cumsum’ function in Matlab was used to get the cumulative sum of how many cells have reoriented at a certain time point, which was then normalized to 1 (depicting ‘all cells’) by dividing by the total number of cells in the dataset.

#### Morphodynamic map analysis

Morphodynamic map analysis was based on the similar analysis in (Yang, Collins, and Meyer 2015). The cell displacement was followed from frame to frame, providing information, where cell plasma membrane was protruding and where it was retracting. In practice, the contour of the cell in each frame of the movie was equidistantly divided into 100 points, called markers. From one frame to another, the pairing of markers was chosen by minimizing the total square distances between markers at time t and t+dt by testing all possible circular shifts of the contour at t+dt. The position of the Nucleus-Golgi axis, calculated from the Golgi and nucleus centroid positions, was plotted on top of the morphodynamic map, showing which way it was pointing. For further analysis and visualization, each column of the obtained morphodynamic maps could be (circularly) shifted, so that the middle marker always represents the Nucleus-Golgi axis, cell trajectory or the x-axis of the image by using Matlab’s ‘circshift’ function.

##### Autocorrelation plot

Autocorrelation data were plotted following the (Gorelik and Gautreau 2014) paper, but adapted from Excel to Matlab.

##### Cross-correlation analysis of traffic flow

Image stacks in tiff format were adjusted using Fiji’s ‘Bleach correction - Histogram matching’ function. Then, a previously described morphodynamic map of cell protrusions was recorded for every cell and a Golgi mask was created using the segmentation algorithm described in the section of ‘Image segmentation’. A line was drawn from every one of 100 points of the cell boundary in the morphomap towards the centroid of the Golgi mask. Using Matlab’s function ‘improfile’, the mean of fluorescence intensity was calculated along every line. Using these calculations another morphomap, depicting the secretion pattern, was drawn and cross-correlation between the two morphomaps was calculated using ‘xcov’ function in Matlab.

### Modelling

See supplementary materials – Model

## Supplemental information

**Supplementary Figure 1.** Feedback routine for moving the microscope stage (field of view) to follow a migrating cell.

**Supplementary Figure 2.** Detailed analysis of Nucleus-Golgi axis movement when a cell is escaping the pattern.

**Supplementary Figure 3.** All morphodynamic maps from control and Nocodazole experiments.

**Supplementary Figure 4.** Examples of RUSH-SPI assays in control and Nocodazole conditions, and all protrusion and trafficking maps from trafficking experiments.

**Supplementary Figure 5.** Biochemical gradient of Cdc42 stabilizes the Golgi complex position and reorients it on an isotropic pattern.

### Supplementary Material Model

Detailed explanation of the minimal physical model.

**Supplementary Movie 1**. RPE1 cell freely moving on a fibronectin covered coverslip followed by a moving microscope stage.

**Supplementary Movie 2**. RPE1 cell “escaping” the pattern.

**Supplementary Movie 3**. RPE1 cell freely moving in 2D in control conditions (a) and with Nocodazole (0.1 μM).

**Supplementary Movie 4**. RPE1 cells in RUSH-SPI assay with labelled Collagen X cargo in control and with Nocodazole, and with labelled Rab6A in control and with Nocodazole.

**Supplementary Movie 5**. RPE1 cells exposed to local optogenetic Cdc42 activation while freely moving, on a round pattern and freely moving with Nocodazole.

**Supplementary Movie 6**. Movement of a synthetic cell recapitulated by the minimal physical model.

