## Supplementary Material with movie legend for "Persistent cell migration emerges from a coupling between protrusion dynamics and polarized trafficking"

**Supplementary Figure 1.**

Feedback routine for moving the microscope stage (field of view) to follow a migrating cell.

(A) First frame ($t_0$) - centroid of a cell (red circle) found from a segmented image; Following frame (t) - new centroid of a cell (blue circle) is found from a taken image at time t, if the two centroids are further than a defined distance, the microscope stage is moved and two centroids are aligned. This routine continues until the end of experiment.

(B) Specific light activation can be included in the feedback routine, where activation pattern is automatically adapted to the shape of the cell and goes along the plasma membrane border with selected pattern thickness.

(C) Morphodynamic map of a representative cell recentered to direction of movement (black).

(D) Average protrusion speed over time (n=17 cells, dashed blue line - sd).

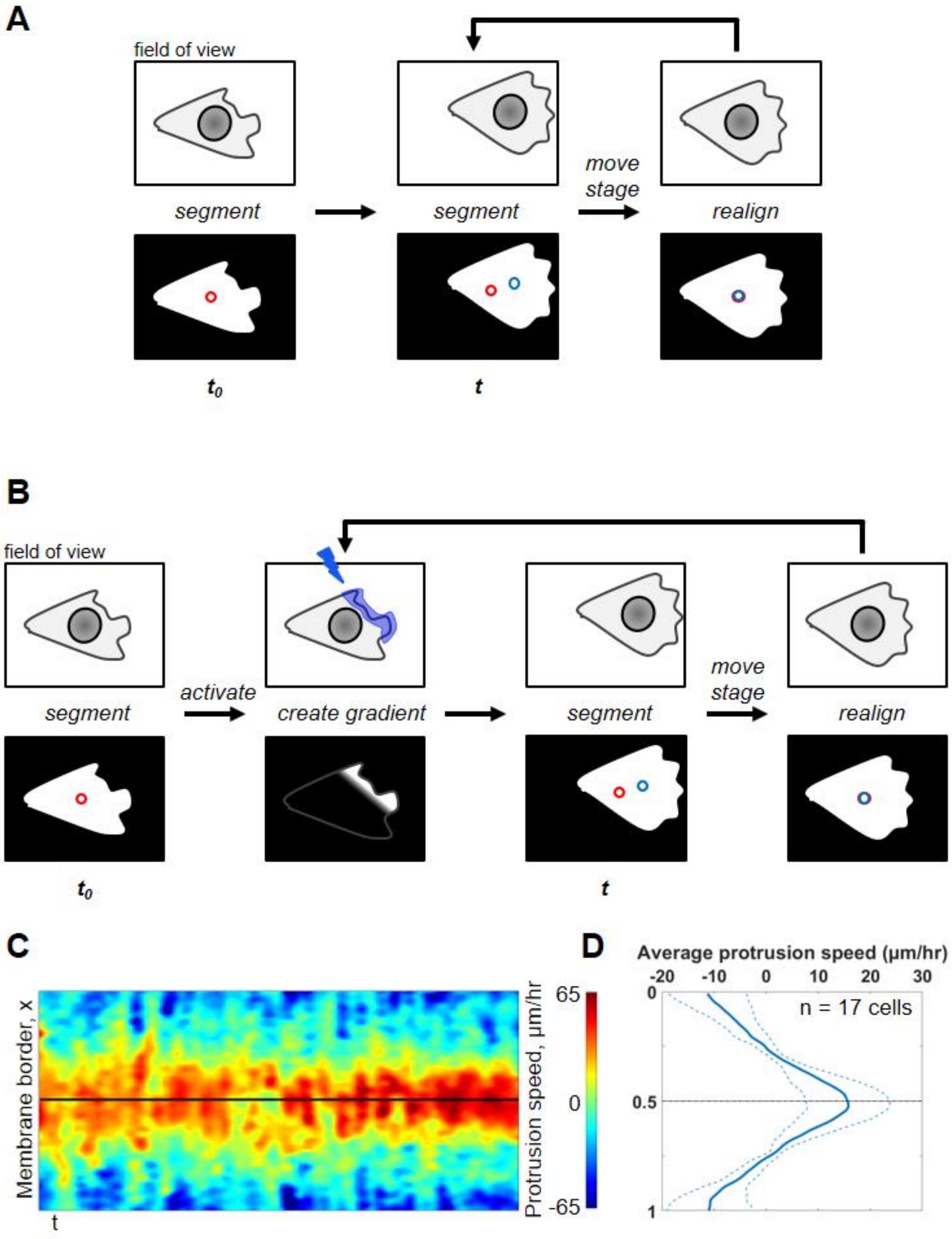

**Supplementary Figure 2.**

Detailed analysis of Nucleus-Golgi axis movement when a cell is escaping the pattern.

(A) Change in position of Nucleus-Golgi axis while moving out of an isotropic pattern (white circle: pattern, a line represents evolution of Nucleus-Golgi axis position in time from blue (start of experiment) to red (end of experiment), movement aligned by the author for visual purposes).

(B) Distance of the nucleus centroid from the center of the pattern. The nucleus starts to move out of the center approximately 1h before the escape. Its movement clearly points to the moment when the cell goes out of the pattern (black arrow) (time resolution – 10 min). Time is normalized for each cell by the time of escape (defined as t=0 here).

(C) Nucleus-Golgi axis position is roughly established at the same time as direction of movement (notice the orange and black arrows pointing to the start of the slope). Evolution of Nucleus-Golgi axis position and cell movement (orange: Direction of movement, defined as Nucleus-center of the pattern axis, black: Nucleus-Golgi axis, the position of both axes are normalized by the direction of movement at the time of "escape" from the pattern (orange curve goes to zero) (n=36 cells), time resolution – 10 min).

A

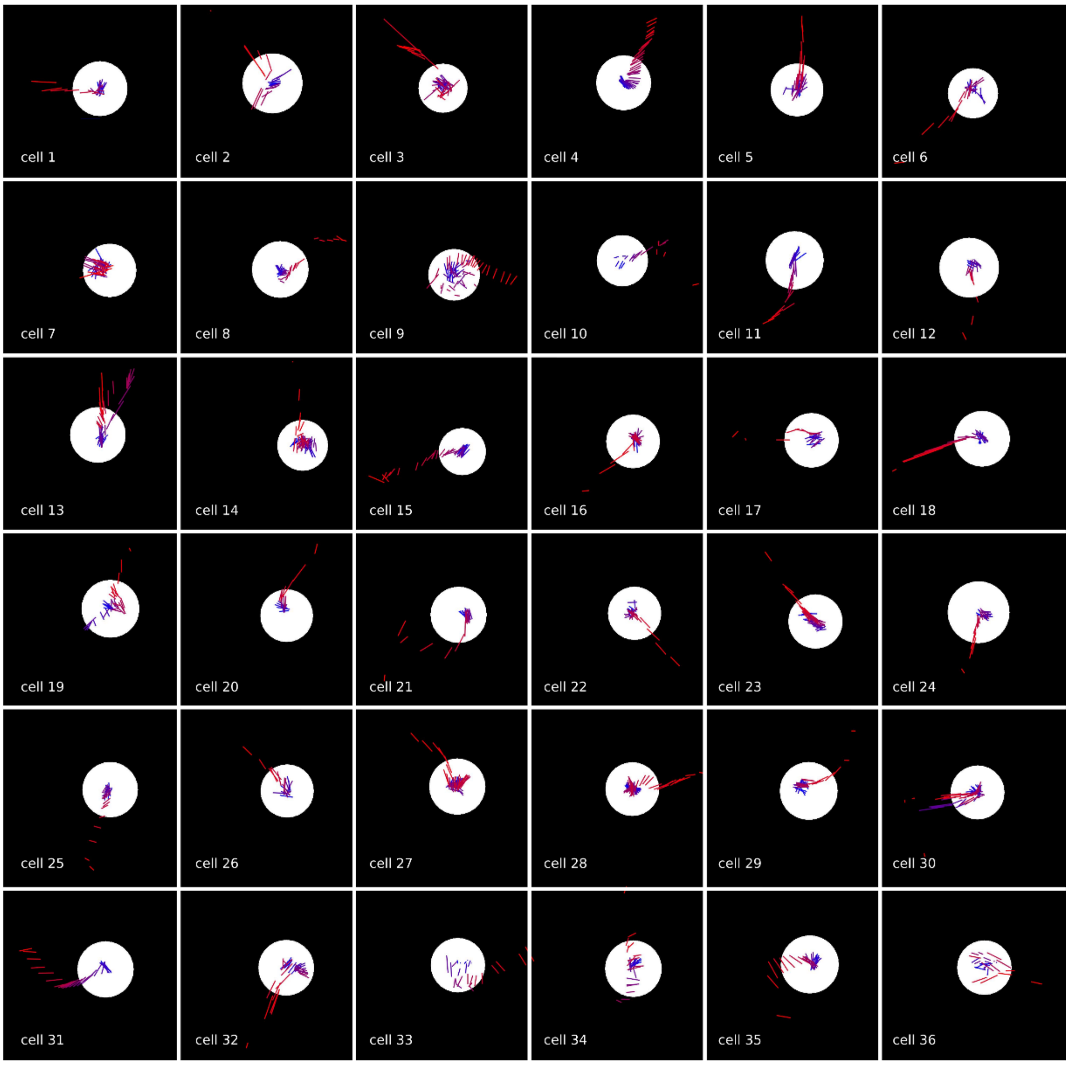

**B**

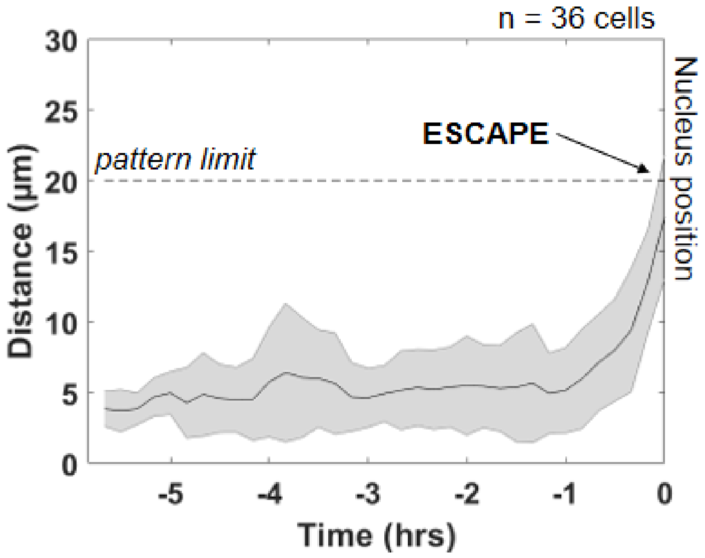

n = 36 cells
Distance (µm)
30
25
20
15
10
5
0
pattern limit
ESCAPE
Nucleus position
-5
-4
-3
-2
-1
0
Time (hrs)


**C**

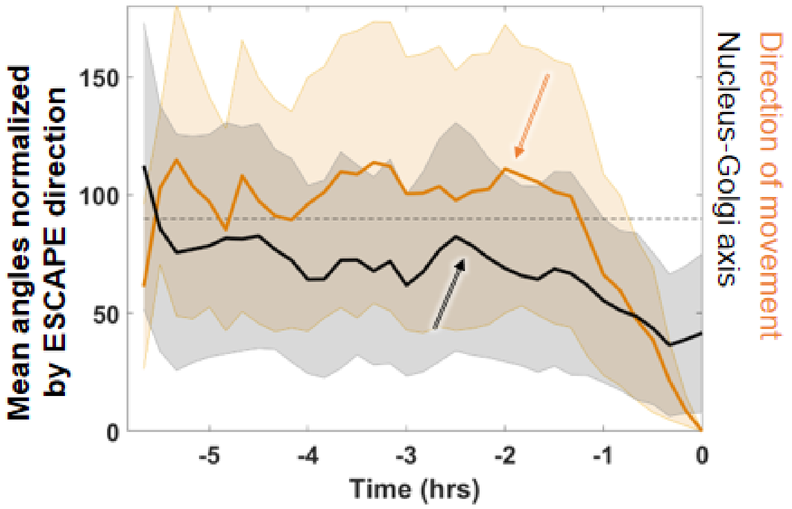

Mean angles normalized by ESCAPE direction
150
100
50
0
Nucleus-Golgi axis
Direction of movement
-5
-4
-3
-2
-1
0
Time (hrs)

**Supplementary Figure 3.**

(A-B) All morphodynamic maps in Ctrl (A) and with NZ (B).

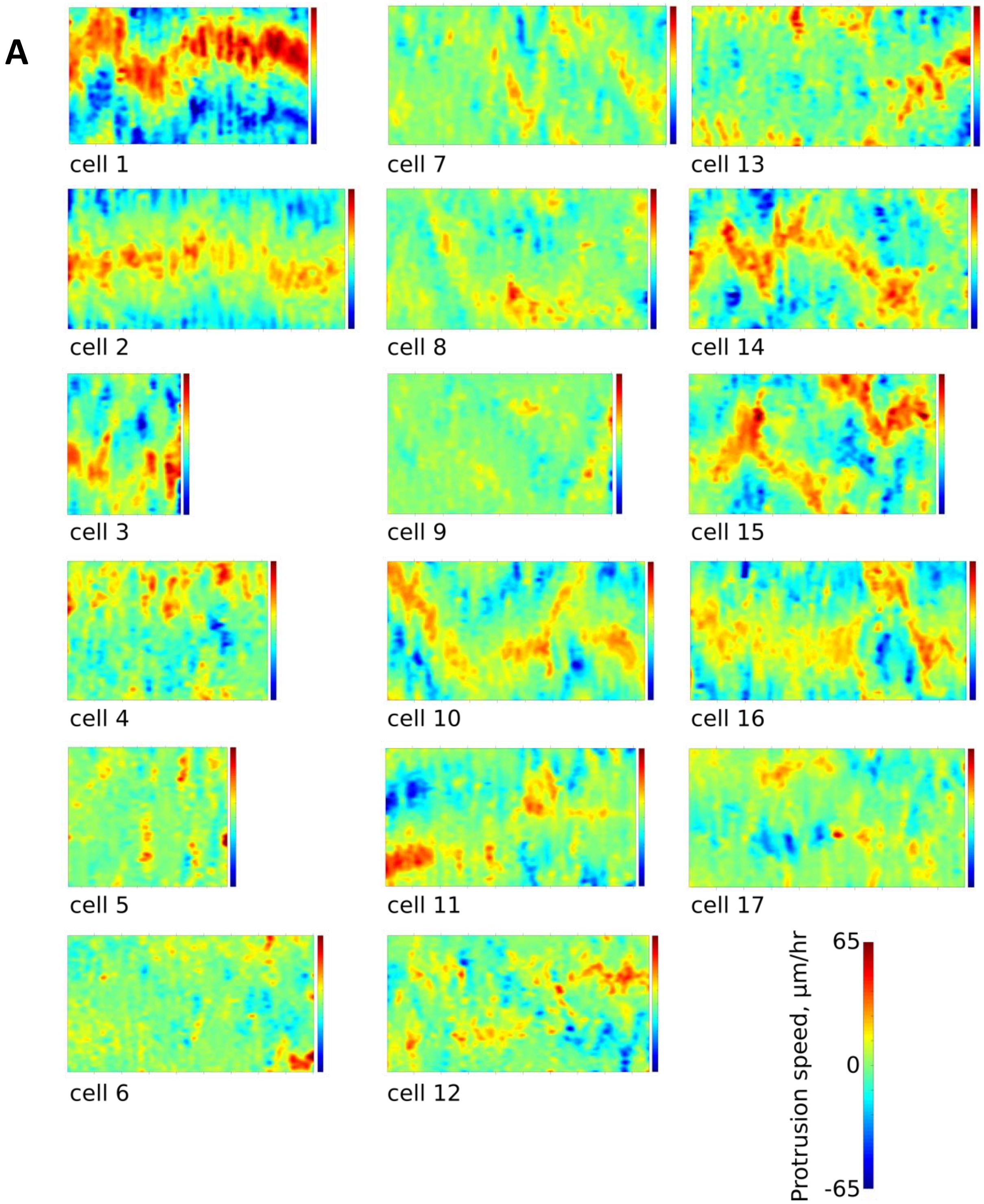

**B**

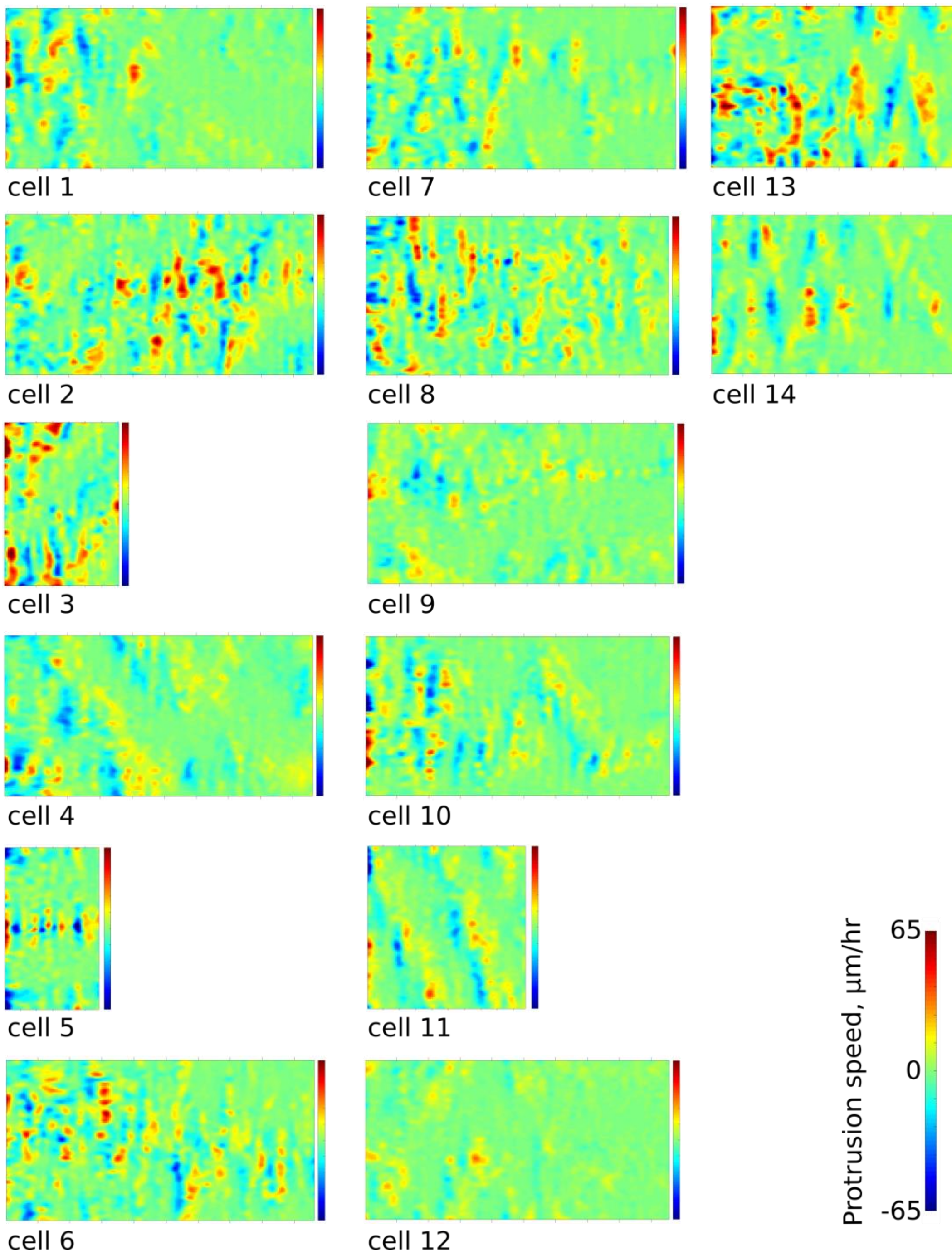

cell 1
cell 7
cell 13
cell 2
cell 8
cell 14
cell 3
cell 9
cell 4
cell 10
cell 5
cell 11
cell 6
cell 12
65
0
-65
Protrusion speed, μm/hr

**Supplementary Figure 4.**

Examples of RUSH-SPI assays in control and Nocodazole conditions, and all protrusion and trafficking maps from trafficking experiments.

(A) Examples of secreted cargo being trafficked towards cellular protrusions in RPE1 cells. Secreted Col10A cargo signature on anti-GFP covered coverslip (green dashed line: protrusions, scale bar - 20 µm).
(B) Examples of cargo secretion in the Nocodazole (0.1 uM) condition in RPE1 cells. Secreted Col10A cargo signature on anti-GFP covered coverslip (scale bar - 20 µm).
(C) All morphodynamic protrusion and trafficking maps in Ctrl condition.
(D) Secretion arrest in presence of GCA. (I-II) The Golgi complex visible in (I) is dispersed upon addition of GCA (35 µM) (II). (III-IV) Biotin (40 µM) is added to release the cargo from the ER and start the RUSH assay (III). (IV) Even after 2 h no noticeable cargo secretion is visible (a mid-section of the cell is shown, where the cargo is trapped) (black label: iRFP-Rab6A in (I) and (II) and Col10A in (III) and (IV), black contour: cell borders, orange: contour of nucleus, scale bar - 20 µm). (V-VI) Post-Golgi vesicle traffic to the protrusion stops in presence of GCA. (V) Post-Golgi vesicles (GFP-Rab6A label) are trafficked towards a forming protrusion before addition of Golgicide A (GCA, 35 µM) (orange arrows point to vesicle accumulation). (VI) Trafficking stops upon addition of GCA (excerpts from the same single-cell movie with constant image brightness parameters, scale bar - 20 µm).

A

B

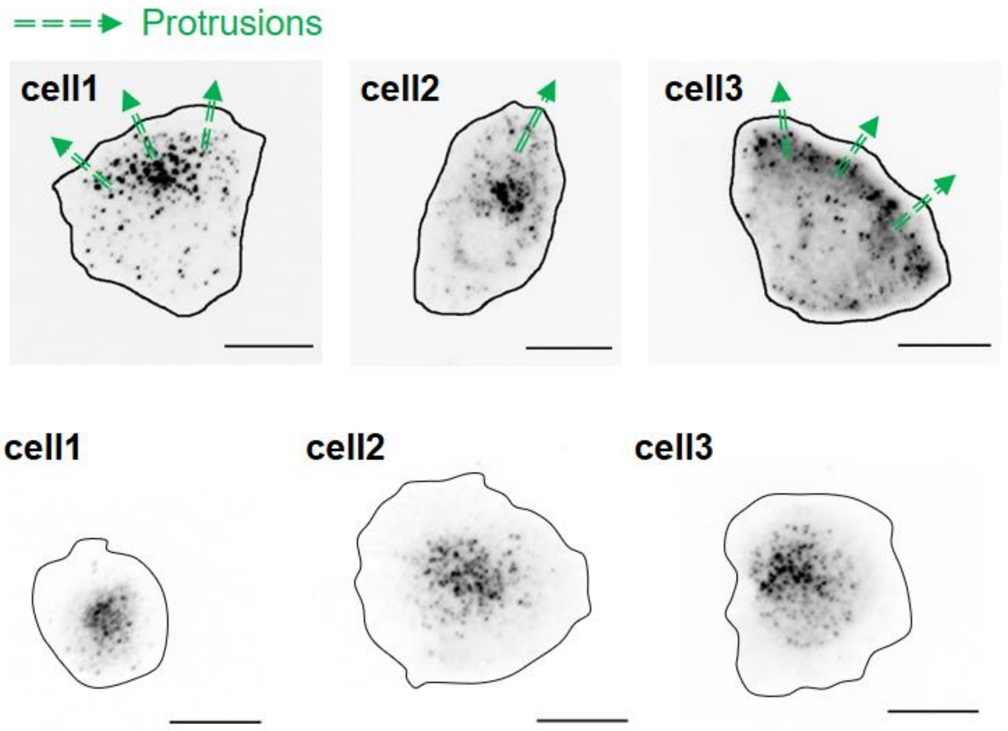

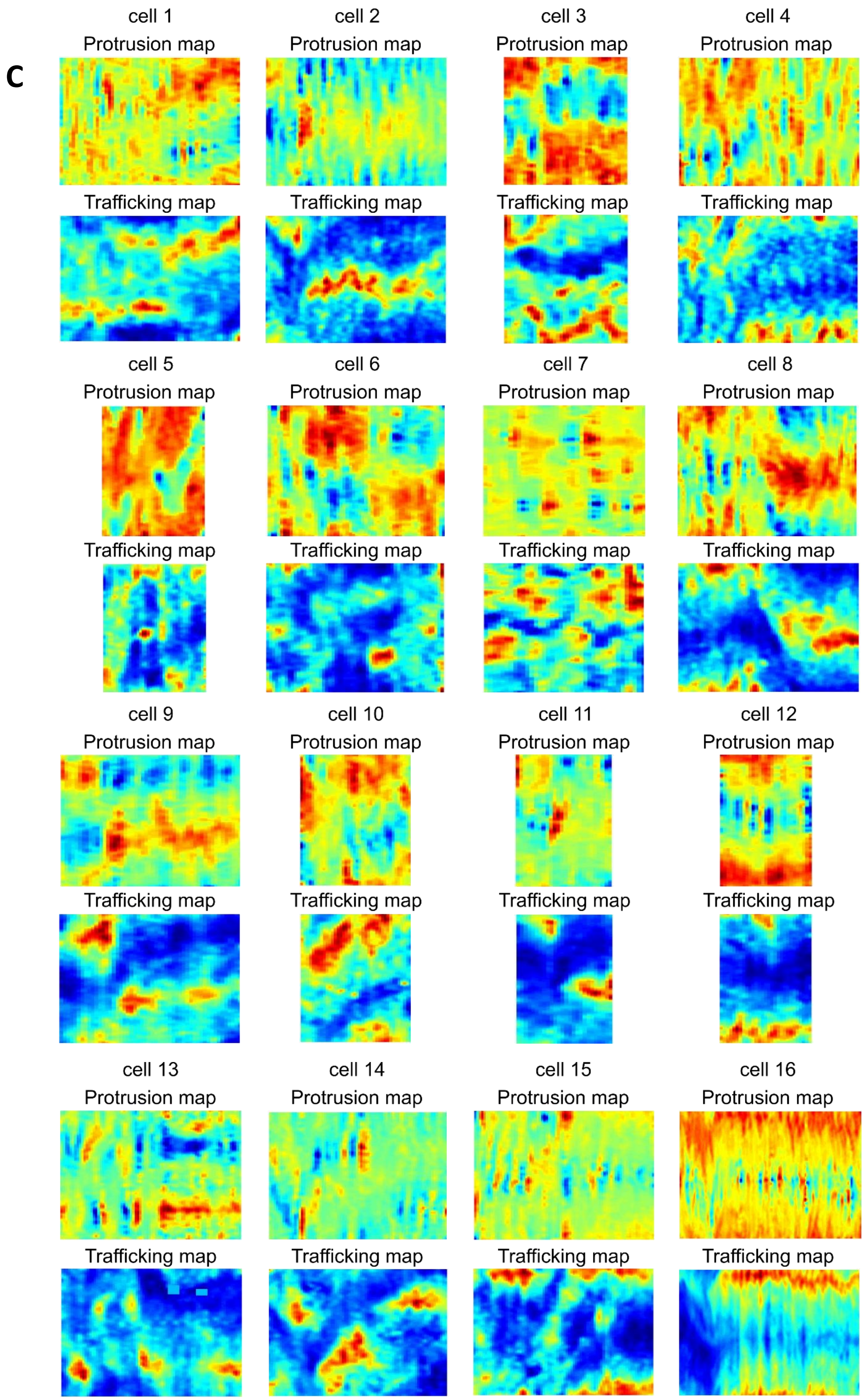
C
cell 1
Protrusion map
Trafficking map
cell 2
Protrusion map
Trafficking map
cell 3
Protrusion map
Trafficking map
cell 4
Protrusion map
Trafficking map
cell 5
Protrusion map
Trafficking map
cell 6
Protrusion map
Trafficking map
cell 7
Protrusion map
Trafficking map
cell 8
Protrusion map
Trafficking map
cell 9
Protrusion map
Trafficking map
cell 10
Protrusion map
Trafficking map
cell 11
Protrusion map
Trafficking map
cell 12
Protrusion map
Trafficking map
cell 13
Protrusion map
Trafficking map
cell 14
Protrusion map
Trafficking map
cell 15
Protrusion map
Trafficking map
cell 16
Protrusion map
Trafficking map

**D**

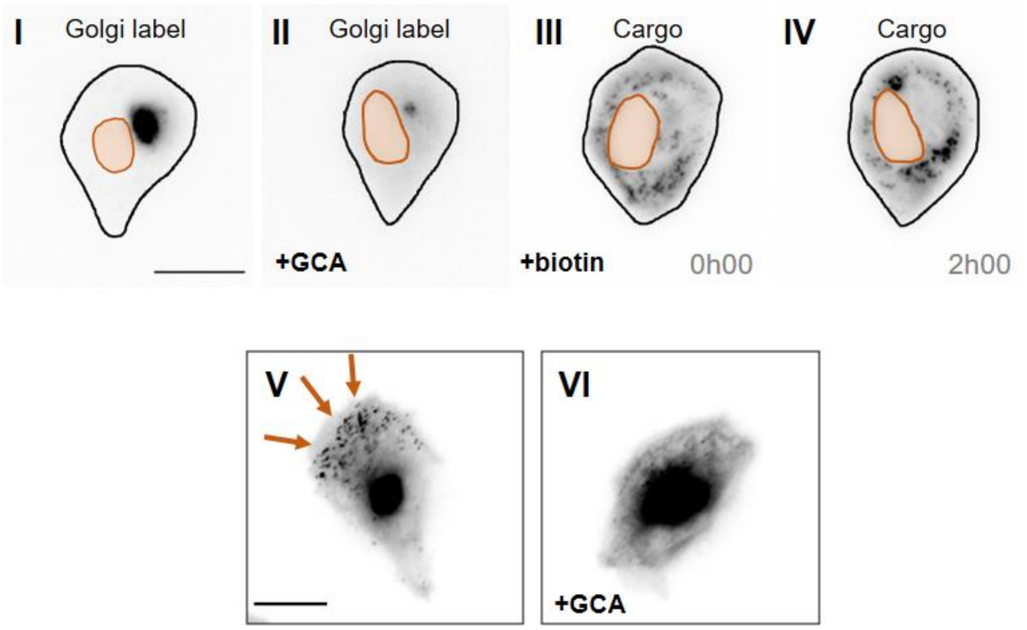

I
Golgi label
II
Golgi label
+GCA
III
Cargo
+biotin
0h00
IV
Cargo
2h00
V
VI
+GCA

**Supplementary Figure 5.**

Biochemical gradient of Cdc42 stabilizes the Golgi complex position and reorients it on an isotropic pattern.

(A) RPE1 cell, following the Nucleus-Golgi axis reorientation toward the optogenetically induced protrusion (yellow: iRFP-Rab6A, blue: optogenetic activation, black dashed line: Nucleus-Golgi axis, blue dashed line: optogenetic activation axis, scale bar – 20 µm).

(B) Optogenetic activation of protrusion formation in front of existing Nucleus-Golgi axis leads to its stabilization in RPE1 cells freely moving on fibronectin covered coverslip (n=19 cell; thin orange lines: single cell data, thick orange line: data average, dashed thick orange lines: standard deviation).

(C) Cumulative graph, showing the ratio of cells, whose Nucleus-Golgi axis reorients in time (reorientation threshold - 30°, black: activation on pattern, light orange: activation control, orange: activation on free cells).

(D) RPE1 cell on a round fibronectin pattern (d=35 µm), following the Nucleus-Golgi axis reorientation toward optogenetic activation of Cdc42 (yellow: iRFP-Rab6A, blue: optogenetic activation, black dashed line: Nucleus-Golgi axis, blue dashed line: optogenetic activation axis, scale bar – 20 µm).

(E-F) Optogenetic activation of protrusion formation 90° away from Nucleus-Golgi axis leads to its reorientation in RPE1 cells on round fibronectin micropatterns (n=16 cells) (E) and is random in control condition (n=13 cells) (F) (thin orange lines: single cell data, thick orange line: data average, dashed thick orange lines: standard deviation).

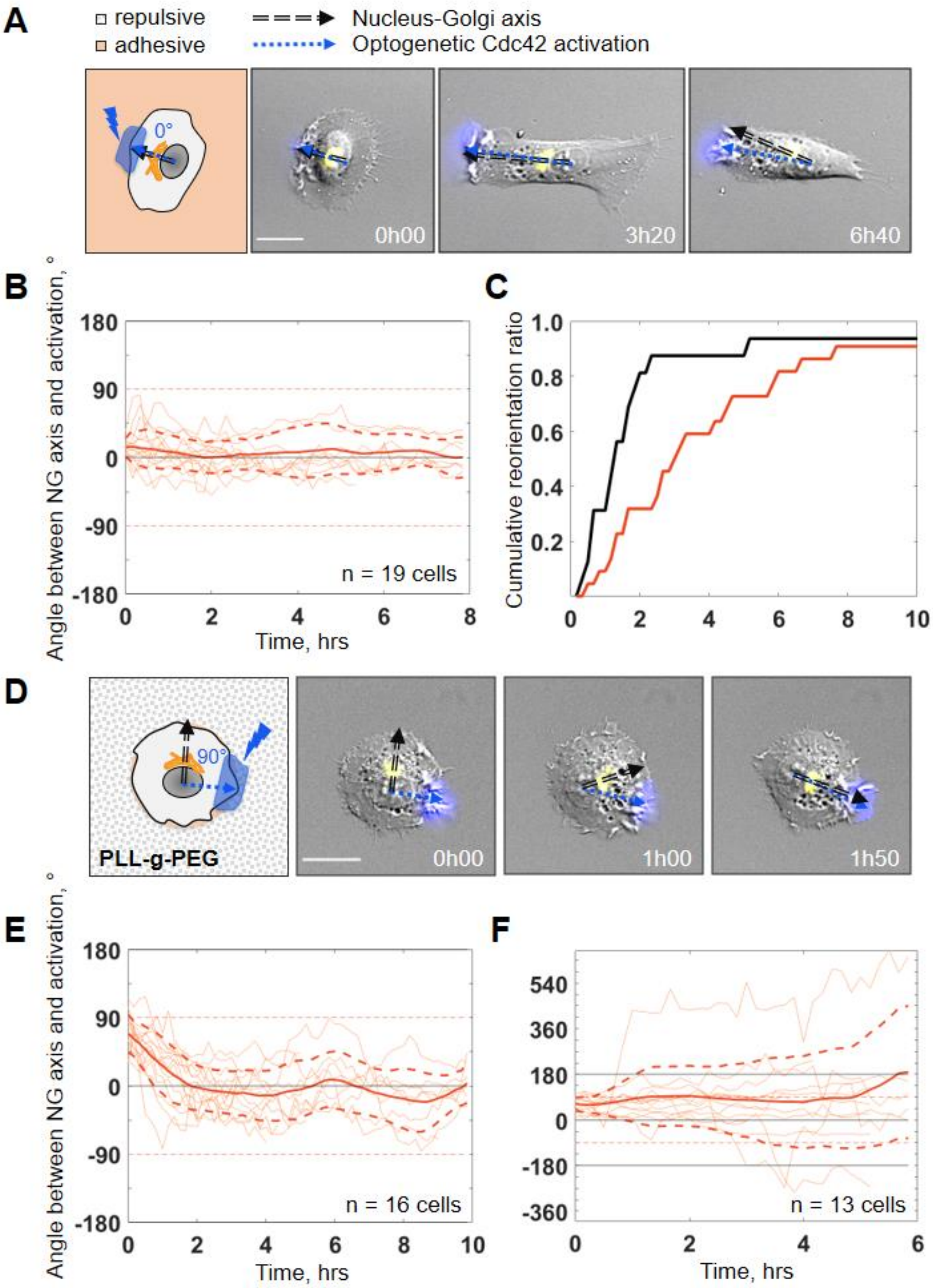

A
repulsive
adhesive
Nucleus-Golgi axis
Optogenetic Cdc42 activation
0°
0h00
3h20
6h40
B
Angle between NG axis and activation, °
180
90
0
-90
-180
n = 19 cells
0
2
4
6
8
Time, hrs
C
Cumulative reorientation ratio
1.0
0.8
0.6
0.4
0.2
0
2
4
6
8
10
D
90°
PLL-g-PEG
0h00
1h00
1h50
E
Angle between NG axis and activation, °
180
90
0
-90
-180
n = 16 cells
0
2
4
6
8
10
Time, hrs
F
540
360
180
0
-180
-360
n = 13 cells
0
2
4
6
Time, hrs

**Supplementary Movie 1**.

RPE1 cell freely moving on a fibronectin (2 µg/mL) covered coverslip followed by a moving microscope stage (cyan: myr-iRFP, yellow: Rab6A, blue: Hoechst 33342, trajectory overlaid in orange, scale bar – 20 µm, time resolution – 5 min.)

**Supplementary Movie 2**.

RPE1 cell "escaping" the pattern (scale bar – 20 µm, time resolution – 10 min).

**Supplementary Movie 3**.

RPE1 cell freely moving on a fibronectin (2 µg/mL) covered coverslip followed by a moving microscope stage in control conditions (a) and with Nocodazole (0.1 µM) (b), scale bar – 20 µm, time resolution – 5 min).

**Supplementary Movie 4**.

(a) RPE1 cell transfected with labelled Collagen X cargo (labelled in black) in control conditions. The cargo is travelling from ER to the Golgi complex and is secreted during a RUSH assay experiment towards a newly forming protrusion (time resolution – 2 min, scale bar - 20 µm).
(b) RPE1 cell transfected with labelled Collagen X cargo (labelled in black) treated with Nocodazole (0.1 µM) (time resolution – 2 min, scale bar - 20 µm).
(c) RPE1 cell with labelled Rab6A (labelled in black) freely moving on a fibronectin (2 µg/mL) covered coverslip in control conditions (black arrow: Nucleus-Golgi axis, cyan arrow: secretion axis, scale bar – 20 µm, time resolution – 5 min).
(d) RPE1 cell with labelled Rab6A (labelled in black) freely moving on a fibronectin (2 µg/mL) covered coverslip treated with Nocodazole (0.1 µM) (scale bar – 20 µm, time resolution – 5 min).

**Supplementary Movie 5**.

RPE1 cells exposed to local optogenetic Cdc42 activation while freely moving, on a round pattern and freely moving with Nocodazole.

(a) RPE1 cell, following the Nucleus-Golgi axis reorientation toward the optogenetically induced protrusion (yellow: iRFP-Rab6A, blue: optogenetic activation, scale bar – 20 µm, time resolution – 10 min).
(b) RPE1 cell on a round fibronectin pattern (d=35 µm), following the Nucleus-Golgi axis reorientation toward optogenetic activation of Cdc42 (yellow: iRFP-Rab6A, blue: optogenetic activation, scale bar – 20 µm, time resolution – 10 min).
(c) RPE1 cell, following the Nucleus-Golgi axis reorientation toward the optogenetically induced protrusion treated with Nocodazole (NZ, 0.1 µm) (yellow: iRFP-Rab6A, blue: optogenetic activation, scale bar – 20 µm, time resolution – 10 min).

**Supplementary Movie 6**.

Movement of a synthetic cell recapitulated by the minimal physical model.
