## Supplementary Material - Model for "Persistent cell migration emerges from a coupling between protrusion dynamics and polarized trafficking"

### Minimal physical model

We present here the minimal physical model of cell migration that implements the two sides of the feedback between an internal polarity axis and protrusion dynamics. It is intended to reproduce and connect our main experimental observables, namely morphodynamic maps and directional persistence of cell trajectories. In this model, we considered the dynamics of the cell edge which ultimately dictates the movement of the cell. Indeed, considering that the cell area is constant (no elastic deformation), the displacement of the cell center of mass is given by the sum of all displacement of points along the edge.

#### Morphodynamic map of a single protrusion

As a first step, we implemented a Matlab© code to simulate realistic morphodynamic maps. Similarly to our data, we considered the movement of 100 markers, $x_i$ with $i \in \{0 \dots 100\}$, equally distributed along the contour of a synthetic cell. To simulate one protrusive event, we implemented numerically a transfer function such that it was similar to the experimental one reported in (Yamao et al. 2015). This transfer function characterizes the edge movement following a point-like activity of Cdc42. To build it, we summed four terms: i) a negative Gaussian function with a relatively large width to account for the initial lateral inhibition, ii) a positive Gaussian function with a relatively small width to account for the actual local protrusive event, iii) a positive Gaussian function with a mean moving forward and backward to account for the traveling wave, and iv) a negative Gaussian function in time centered on 0 to account for the central long-lasting inhibition. All those terms were combined with exponentially decaying function of time to account for the temporal "fading". This transfer function was further convolved with a synthetic Cdc42 signal to simulate a realistic protrusion. We assumed that Cdc42 activity for a single protrusion could be described by a "signaling puff" in space and time, made using a product between a Gaussian function (lateral extension of the puff) and an exponentially decaying function of time (duration of the puff). The resulting morphodynamic map then represents the edge dynamics following a single protrusive event (note that what we call a single protrusive event could be the outcome of several protrusions sustained by another feedback - for example between actin and Cdc42 or Rac1 activity). The duration of Cdc42 activity was chosen such that the duration of the protrusion event was similar to the one observed in our data. For that, we considered the experimental morphodynamic maps under Nocodazole treatment to better isolate “unique” rounds of localized protrusive activity. Experimental protrusions are extended over c.a. 10 frames, thus 50 minutes. We used this number such that our simulation time frame matches the experimental one. The “single” protrusion morphodynamic map (thus corresponding to a typical round of localized protrusive activity) was then normalized such that the integral (sum over space and time) was null (zero mean) to ensure a constant cell area. This morphodynamic map is shown in **Figure 6b** of the main manuscript.

#### First side of the feedback: effect of the polarity axis on protrusions

Next, we simulated a full morphodynamic map by introducing the random appearance of protrusion over time. We took a total duration of the simulation of 1000 frames, $t_{\text{tot}} = 1000$, and the frequency at which protrusions happen was characterized by a probability of appearance per unit of time $P_{\text{rand}} = 0.5$. The **Figure 1** illustrates morphodynamic maps for different values of $P$. If for a given time a protrusion event happened, the position of this protrusion event along the contour was chosen either randomly (uniform distribution over all markers of the contour) or in the direction of the polarity axis (see below), given by the position of a specific marker $c_{\text{axis}}$ that corresponds to the intersection of the internal polarity axis with the cell contour (whose dynamics are described in the next paragraph). The choice of a polarized protrusion was characterized by the probability $P_{\text{polarized}}$ (probability of a randomly placed protrusion is then $1 - P_{\text{polarized}}$), which quantifies the strength of the feedback between the polarity axis and biased initiation of protrusions. When $P_{\text{polarized}} = 0$, protrusions are happening randomly along the contour, and when $P_{\text{polarized}} = 1$ protrusions are always happening in front of the polarity axis (see **Figure 2** for an example). To avoid a non-realistic perfectly straight movement, we introduced a low level of noise for polarized protrusions, namely the position of protrusion was drawn from a normally distributed function whose mean is $c_{\text{axis}}$ and standard deviation is $\sigma_c = 5$ (1/20 of the cell contour). Note that our main result does not depend on the exact value of $\sigma_c$. Indeed, this parameter plays a role only on the asymptotic value of the autocorrelation of direction when $P_{\text{polarized}} \sim 1$, but not on the actual characteristic decay time of the autocorrelation function (which characterizes the persistence time, our main observable).

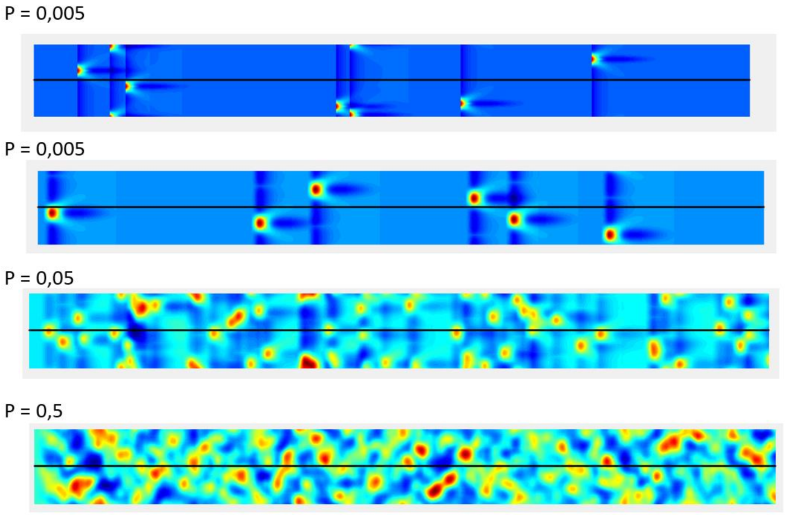


***Figure 1: Example of morphodynamic maps without any feedback for varying frequencies of protrusions.*** *The x-axis is time (1000 frames) and the y-axis is the contour along the cell,* $x_i$ , ; $i \in \{0 \dots 100\}$*. The color code represents membrane speed in arbitrary units, from deep blue (retraction) to deep red (protrusion).*

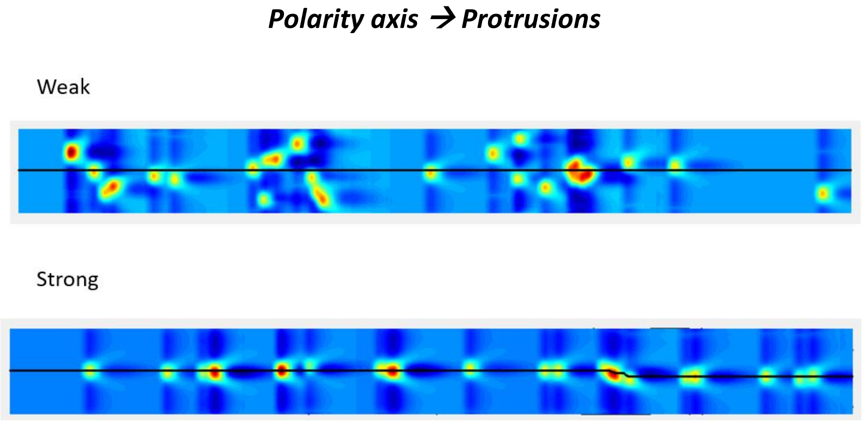


***Figure 2: Example of morphodynamic maps with two extreme values of the feedback between polarity axis and protrusions****. The direction of the polarity axis is depicted by the central black line which corresponds to the position of $c_{axis}$ over time. In the top figure, there is a weak tendency to form protrusion in the direction of the polarity axis. In the bottom figure, there is a strong tendency to form protrusion in the direction of the polarity axis. The x axis is time (1000 frames) and the y axis is the contour along the cell, $x_i$ ; $i \in \{0 \dots 100\}$. The color code represents membrane speed in arbitrary units, from deep blue (retraction) to deep red (protrusion).*

#### Natural dynamics of the polarity axis

As mentioned above, we also needed to explicitly model the evolution of the internal polarity axis, specified by the position $c_{\text{axis}}$, which can be thought as the direction of the polarized secretion or of the Nucleus-Golgi axis. Indeed, in the context of our model, this polarity axis should be taken as the effective bias of either Golgi positioning or polarized secretion on the protrusive activity (both of them correlate, see our experimental data). In absence of any feedback of protrusions on the polarity axis, we could assume that this polarity axis remains at the same position. However, under this assumption, if we plug-in the second feedback (biased protrusions) the cell would move straight, which would not be realistic. We, thus, modeled a "natural" movement of the polarity axis that would mimic the random evolution of the polarity axis when no protrusions affect it. The closest experimental data to infer this dynamic is found for cells plated on a round pattern (see **Supplementary Figure 5f**). In that situation, the polarity axis moves as a correlated random walk whose span reaches 360 degrees in about 4 hours. We simulated such dynamics by assuming that the instantaneous speed of the polarity axis was set by the derivative of a smoothed random function whose values over time are given by a random number between 0 and 10 times the number of markers. The overall natural evolution of the polarity axis, $c_{\text{axis}}^{\text{natural}}$, is depicted in **Figure 3** for a timescale comparable to experiments and in **Figure 4** for the total duration of the simulation. The actual model we chose for the natural evolution of the polarity axis is arbitrary, nonetheless the details do not matter for the outcome of our simulation. The only important effect is that this polarity axis gets reoriented randomly in about 4 hours.

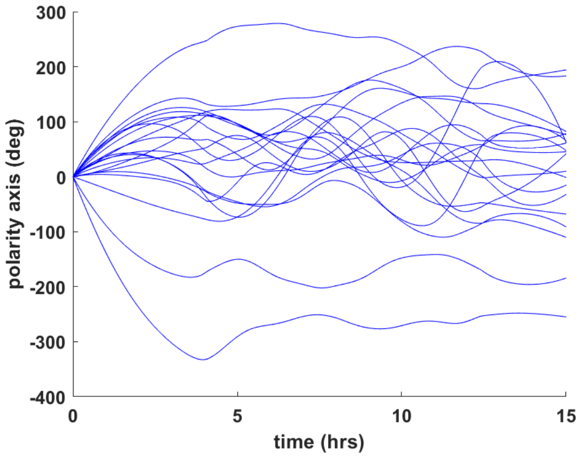


***Figure 3: Natural evolution of the Polarity axis in the simulation for 150 frames** (~15 hours) for 20 repetitions.*

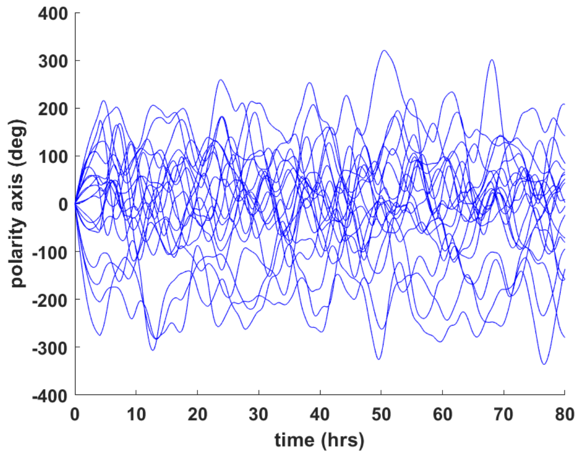


***Figure 4: Natural evolution of the Polarity axis in the simulation for 1000 frames** (~80 hours) for 20 repetitions.*

#### Second side of the feedback: effect of protrusions on the polarity axis

To model the other side of the feedback, namely the reorientation of the polarity axis by protrusions, we assumed that an effective force was attracting the polarity axis towards protrusions. This force was computed as the sum of membrane speed over all markers, $F_{\text{prot}} = \sum_{x_i=0}^{100} v_{x_i}$. The underlying assumption is that every protruding portion of the cell pulls on the polarity axis with a force proportional to the protrusion speed and does this independently of the respective positions of protrusions with regards to the polarity axis. This effective force is competing with the force corresponding to the natural evolution of the Polarity axis, , $F_{\text{basal}} = d/dt\left(c_{\text{axis}}^{\text{natural}}\right)$. The strength of the feedback was implemented by introducing a linear combination between these two forces characterized by a value $\kappa$: $d/dt\,(c_{\text{axis}}) = \kappa F_{\text{prot}} + (1-\kappa)F_{\text{basal}}$. When $\kappa = 0$, the polarity axis follows its natural evolution. When $\kappa = 1$, the polarity axis follows the protrusions. This parameter $\kappa$ can be expressed as the relative contribution of the effective force towards protrusions, $\kappa = (F_{\text{tot}} - F_{\text{basal}})/\left(F_{prot} - F_{\text{basal}}\right)$, where $F_{\text{tot}}$ is the total force acting on the polarity axis. The **Figure 5** shows an example of morphodynamic maps with two values of the strength of the feedback. As above, the rule chosen to implement the feedback is arbitrary (we could have taken a metric such that protrusions close to the polarity axis matter more than the distant ones), but our goal was to implement a minimal model integrating the feedback.

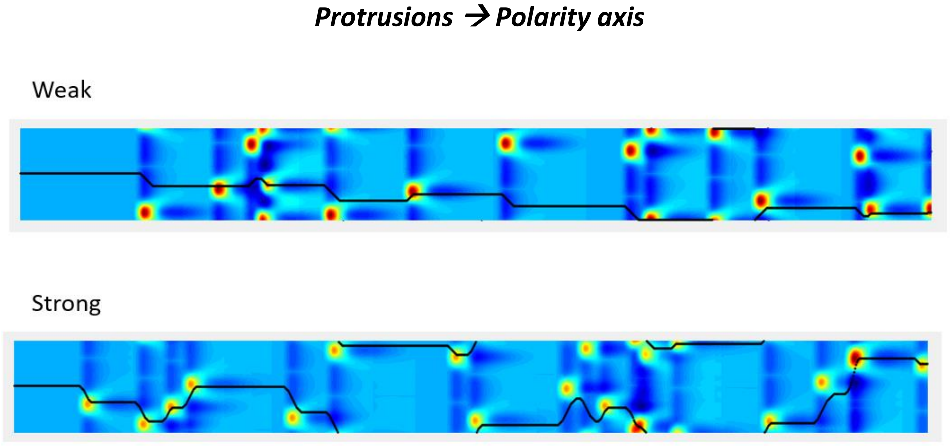


***Figure 5: Example of morphodynamic maps with two extreme values of the feedback between protrusions and polarity axis.*** *The direction of the Polarity axis is depicted by the black line which corresponds to the position of* $c_{axis}$ *over time. In the top figure, there is a weak tendency of the Polarity axis to follow protrusions, whereas in the bottom figure, there is a strong tendency of the Polarity axis to align with every protrusion. The x-axis is time (1000 frames) and the y-axis is the contour along the cell,* $x_i$ ; $i \in \{0 \dots 100\}$. *The color code represents membrane speed in arbitrary units, from deep blue (retraction) to deep red (protrusion).*

#### Full model with the two-sided feedback

Altogether, the two sides of the feedback can be implemented in our simulation to produce realistic morphodynamic maps, see **Figure 6** for an example with a low frequency of protrusions and **Figure 7** for an example with realistic protrusion frequency. These morphodynamic maps were then translated into cell trajectories, by noticing that the instantaneous movement of the cell centroid is simply the sum of all the velocities of the markers along the cell contour. These trajectories were analyzed in the same way as the experimental ones, to obtain the autocorrelation of direction and, then, the characteristic decay time of this function was calculated (persistence time). By varying the strength of the two sides of the feedback, we were able to produce the phase diagram presented in **Figure 6g** of the main manuscript. For visualization purposes, we also used the morphodynamic maps to produce synthetic movies of migrating cells (see **Supplementary movie 6**). For this purpose, we inverted the process of morphodynamic map quantification and used a morphodynamic map to evolve an elastic contour over time.

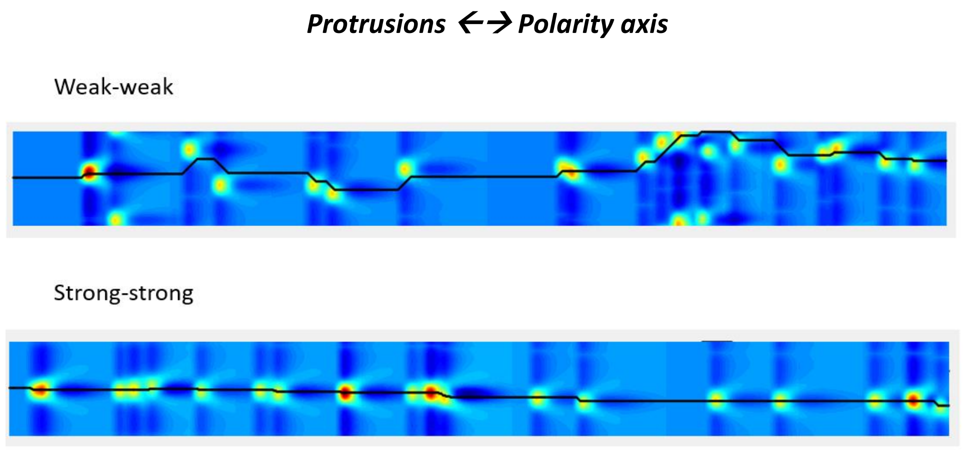


*__Figure 6: Example of morphodynamic maps with the two sides of the feedbacks in action.__ The direction of the Polarity axis is depicted by the black line, which corresponds to the position of* $c_{axis}$ *over time. The top figure corresponds to weak feedback, and the bottom figure to strong feedback. The x-axis is time (1000 frames) and the y-axis is the contour along the cell,* $x_i$ ; $i \in \{0 \dots 100\}$*. The color code represents membrane speed in arbitrary units, from deep blue (retraction) to deep red (protrusion).*

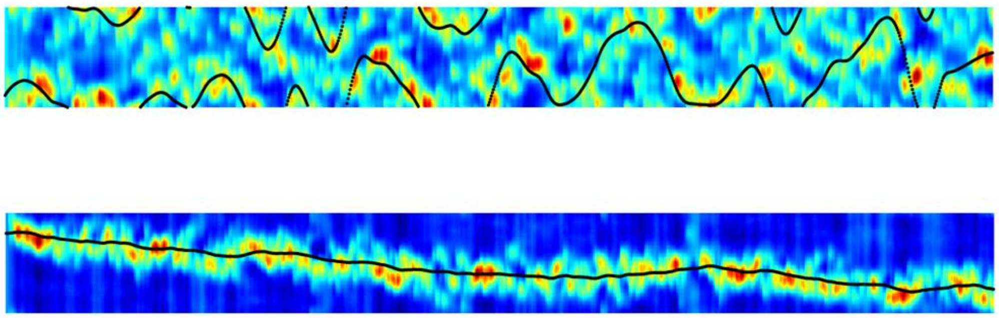

*__Figure 7: Example of morphodynamic maps with the two sides of the feedbacks in action, for realistic frequencies of protrusion per unit of time__. The direction of the Polarity axis is depicted by the black line which corresponds to the position of* $c_{axis}$ *over time. the top figure corresponds to intermediate feedback (*$\kappa = 0.5$; $P_{\text{polarized}} = 0.5$*) that still leads to random migration. The bottom figure corresponds to strong feedback (*$\kappa = 1$; $P_{\text{polarized}} = 1$*) that leads to very persistent migration. The x-axis is time (1000 frames) and the y-axis is the contour along the cell,* $x_i$ ; $i \in \{0 \dots 100\}$*. The color code represents membrane speed in arbitrary units, from deep blue (retraction) to deep red (protrusion).*

#### Protrusion unicity index

In addition to the mapping of persistence time as a function of the strength of the two sides of the feedback, we developed two other independent measures aimed at quantifying cell polarity. First, to quantitatively assess the polarity of cells in terms of front-to-back polarity, we wanted to compute a number, characterizing the number of simultaneous protrusions. Indeed, cells presenting a single protrusion over time are monopolar and thus well polarized, whereas cells presenting several protruding fronts are multipolar and, thus, are lacking a single polarity axis. We introduced a Protrusion unicity index as the inverse average number of simultaneous protrusions to quantify this aspect of cell polarity. To compute this index, we used a sliding window of 10 frames (50 minutes) applied on the morphodynamic map and we segmented protrusions based on a threshold (70\% of maximal protrusion speed). The number of competing protrusions, $N_p$, was then obtained as the number of non-connected segmented objects and this number was averaged over the whole duration of the simulation (1000 frames) and over

20 realizations. The Protrusion unicity index for a given value of $\kappa$ and $P_{\text{polarized}}$ was taken as $1/N_p$. This index is equal to one when a single protrusion is present, and goes to zero as more and more protrusions are competing. As seen from the **Figure 8**, the main parameter dictating this unicity index is $P_{\text{polarized}}$. This is easily understood: no matter if the polarity axis follows protrusions or not, if $P_{\text{polarized}}$ is high, there will always be a single protrusive activity in front of the polarity axis.

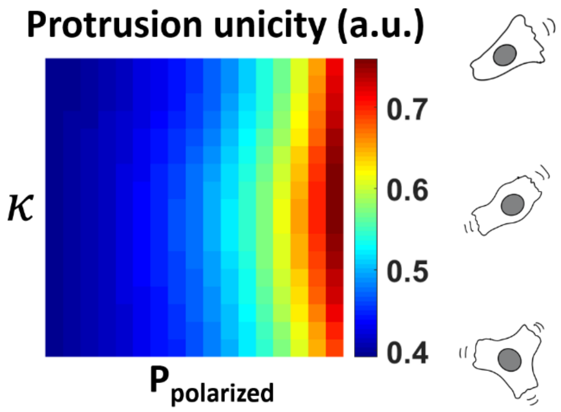


***Figure 8: Protrusion unicity index.*** *The three sketches of a migrating cell next to the scale represent the different amount of protrusions along the scale.*

#### Alignment index

To quantitatively assess the polarity of cells in terms of the alignment of their polarity axis with direction of movement, we wanted to compute how well the polarity axis aligns with cell movement. We, thus, introduced an Alignment index, as the standard deviation of the angle between the polarity axis and instantaneous direction of movement. In terms of circular statistics, this standard deviation is called the angular dispersion and is defined as:

$$r = \sqrt{\left(\frac{1}{n}\sum_{i=1}^{n}\sin\theta_i\right)^2 + \left(\frac{1}{n}\sum_{i=1}^{n}\cos\theta_i\right)^2}$$

The angular dispersion varies between 0 (uniform dispersion) and 1 (perfect alignment). As seen from the **Figure 9**, the Alignment index depends both on $\kappa$ and $P_{\text{polarized}}$. Indeed, when $P_{\text{polarized}}$ is close to zero, even if $\kappa = 1$ protrusions happen randomly all the time and the polarity axis does not have time to follow them, thus leading to a low value of alignment.

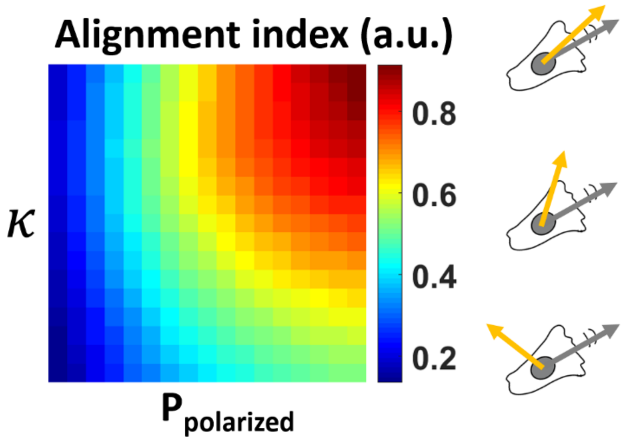


***Figure 9: Alignment index.*** *The three sketches of a migrating cell next to the scale represent the different angles of alignment between polarity axis and direction of movement along the scale.*
